# Structure of the *Lysinibacillus sphaericus* Tpp49Aa1 pesticidal protein elucidated from natural crystals using MHz-SFX

**DOI:** 10.1101/2022.01.14.476343

**Authors:** Lainey J. Williamson, Marina Galchenkova, Hannah L. Best, Richard J. Bean, Anna Munke, Salah Awel, Gisel Pena, Juraj Knoska, Robin Schubert, Katerina Doerner, Hyun-Woo Park, Dennis K. Bideshi, Alessandra Henkel, Viviane Kremling, Bjarne Klopprogge, Emyr Lloyd-Evans, Mark Young, Joana Valerio, Marco Kloos, Marcin Sikorski, Grant Mills, Johan Bielecki, Henry Kirkwood, Chan Kim, Raphael de Wijn, Kristina Lorenzen, P. Lourdu Xavier, Aida Rahmani, Luca Gelisio, Oleksandr Yefanov, Adrian P. Mancuso, Brian Federici, Henry N. Chapman, Neil Crickmore, Pierre J. Rizkallah, Colin Berry, Dominik Oberthür

**Affiliations:** School of Biosciences, Cardiff University, UK; Center for Free Electron Laser Science CFEL, Deutsches Elektronen-Synchrotron DESY, Notkestr. 85, 22607 Hamburg, Germany; European XFEL GmbH, Schenefeld, Germany; Department of Biological Sciences, California Baptist University, USA; Max-Planck Institute for the Structure and Dynamics of Matter, 22761 Hamburg, Germany; Department of Chemistry and Physics, La Trobe Institute for Molecular Science, La Trobe University, Melbourne, Victoria 3086, Australia; Department of Entomology and Institute for Integrative Genome Biology, University of California, USA; Centre for Ultrafast Imaging, Universität Hamburg, Hamburg, Germany; Department of Physics, Universität Hamburg, Hamburg, Germany; School of Life Sciences, University of Sussex, Falmer, UK; School of Medicine, Cardiff University, UK

**Keywords:** XFEL, SFX, *Lysinibacillus sphaericus*, Pesticidal protein, Tpp49Aa1, Cry48Aa1

## Abstract

Tpp49Aa1 from *Lysinibacillus sphaericus* is a Toxin_10 family protein that – in combination with Cry48Aa1, a 3-domain crystal protein - has potent mosquitocidal activity, specifically against *Culex quinquefasciatus* mosquitoes. MHz serial femtosecond crystallography at a nano-focused X-ray free electron laser, allowed rapid and high-quality data collection to determine the Tpp49Aa1 structure at 1.62 Å resolution from native nanocrystals. This revealed the packing of Tpp49Aa1 within these nanocrystals, isolated from sporulated bacteria, as a homodimer with a large intermolecular interface, shedding light on natural crystallization. Complementary experiments conducted at varied pH also enabled investigations of the early structural events leading up to the dissolution of natural Tpp49Aa1 crystals. Using modelling, we propose a potential interaction between Tpp49Aa1 and Cry48Aa1 that may play a role in their codependency and broaden our understanding of this two-component system. We expand the known target range, demonstrating Tpp49Aa1/Cry48Aa1 susceptibility of larvae from *Anopheles stephensi, Aedes albopictus* and *Culex tarsalis* – substantially increasing the potential use of this toxin pair in mosquito control. Further functional insights are gained using *Culex* cell lines to characterise cellular models for future investigations into Cry48Aa1/Tpp49Aa1 mechanism of action and to demonstrate transient detrimental effects of individual toxin components.

**Significance Statement:** The Tpp49Aa1/Cry48Aa1 protein pair kills mosquito larvae. Innovative use of nano-focused X-ray free electron laser to match the size of natural Tpp49Aa1 nanocrystals and the highest beam intensity available in any XFEL for high-throughput data collection, allowed structural resolution to 1.62 Å.

Tpp proteins show a range of interactions with different partners to elicit toxicity. To gain insight into Tpp49Aa1, its interaction with Cry48Aa1 was modelled. We also establish cell-based assays of Tpp49Aa1/Cry48Aa1 activity.

We expand the known target range to include three more mosquito species: *Anopheles stephensi, Aedes albopictus* and *Culex tarsalis*. This study will underpin future Tpp mode of action investigations and aid insecticide optimization against mosquito vectors of emerging diseases such as West Nile Virus and malaria.

## 1. Introduction

Pesticidal proteins produced by *Bacillus thuringiensis* and *Lysinibacillus sphaericus* constitute the major factors in bioinsecticides and transgenic crops (1). The mosquitocidal Tpp1Aa2/Tpp2Aa2 (formerly BinA2/BinB2) pair is produced by highly pathogenic strains of *L. sphaericus* (2, 3) and has been commercially applied in the field to control mosquito vectors of human disease. Despite its success, cases of resistance have been identified (4, 5). To address this, *L. sphaericus* isolates exhibiting toxicity against Tpp1Aa2/Tpp2Aa2-resistant *Culex quinquefasciatus* mosquito larvae were screened (6, 7), leading to the identification of the Cry48Aa1/Tpp49Aa1 pair (7–9).

The Cry48Aa/Tpp49Aa pair is produced by some *L. sphaericus* strains (including NHA15b and IAB59), and is composed of Cry48Aa, a 135 kDa protein belonging to the 3-domain family of Crystal (Cry) proteins, and Tpp49Aa (formerly Cry49Aa), a 53 kDa protein belonging to the family of Toxin_10 pesticidal proteins (Tpp) (8, 10). Both proteins are deposited in the form of natural, parasporal crystals in association with the bacterial spore. Mosquito toxicity bioassays have revealed that both components in combination are required for activity against *C. quinquefasciatus* larvae, with optimal activity at a ∼1:1 molar ratio (8, 9). The Cry48Aa1/Tpp49Aa1 toxin pair does not show toxicity to any other insect species investigated, including the Coleopteran, *Anthonomus grandis*, the Lepidoptera, *Anticarsia gemmatalis, Spodoptera frugiperda* and *Plutella xylostella*, and the Diptera, *Aedes aegypti* and *Anopheles gambiae* (9). When administered in purified protein form, the potency of the Cry48Aa1/Tpp49Aa1 pair against *C. quinquefasciatus* (LC_50_: 15.9 and 6.3 ng/ml, respectively for the co-administered proteins) (8) is comparable to that of Tpp1Aa2/Tpp2Aa2 (LC_50_: ∼30 ng/ml each) (11). Hence, Cry48Aa1/Tpp49Aa1 could be applied in the development of new pesticidal agents aimed at overcoming Tpp1Aa2/Tpp2Aa2 resistance.

In common with other pesticidal proteins, the mode of action of the Cry48Aa1/Tpp49Aa1 toxin pair begins with ingestion of crystal inclusion protoxins, followed by solubilization and proteolytic cleavage in the alkaline environment of the gut (8). Following activation, both Cry48Aa1 and Tpp49Aa1 bind specifically and with high affinity to *C. quinquefasciatus* brush border membrane (12). Although functional receptors are yet to be elucidated, several classes of Cry48Aa1/Tpp49Aa1 binding proteins have been identified, including maltases, aminopeptidases, alkaline phosphatases, and metalloproteases (13). Following membrane interaction, subsequent cytopathological effects including cytoplasmic and mitochondrial vacuolation, endoplasmic reticulum breakdown, and microvillus disruption are seen (14), culminating in insect death.

Dot blot assays show Cry48Aa1 and Tpp49Aa1 are able to form a complex (12). Moreover, competition assays have indicated that the N-terminal region of Tpp49Aa1, residues Asn49 – Ser148, is responsible for binding Cry48Aa1, while the C-terminal region, residues Ser349 – Asn464, is involved in membrane interaction (12). Considering the combined role of Tpp49Aa1 in both interaction with Cry48Aa1 and the target cellular membrane, as well as studies indicating a significant loss of larvicidal activity from Tpp49Aa1 truncated fragments, it has been suggested that the region of Tpp49Aa1 required for activity corresponds to the protein fragment remaining after proteolytic activation after residue Phe48 (9, 12). Mutagenesis studies targeting Tpp49Aa1 cysteine residues 70, 91, 183, and 258, have indicated the functional importance of the latter three, which are required for full larvicidal activity against *Culex* larvae, as well as Cry48Aa1 binding (15). Although homology modelling has been applied to produce structural models of both Cry48Aa1 and Tpp49Aa1 (9, 16), their structures are yet to be experimentally resolved.

The development of X-ray free electron lasers (XFELs) has given rise to a new approach in protein crystallography, known as serial femtosecond crystallography (SFX). SFX introduces a stream of crystals into an XFEL beam, where the delivery of intense X-ray pulses of several femtoseconds duration enables diffraction data to be collected at high exposures in a serial fashion before structural information is lost due to radiation damage (17–19). As such, diffraction data are not limited by the small crystal size of natural crystals. In the field of pesticidal proteins, SFX has previously been applied to solve the structures of the Cyt1Aa, Cry3Aa and Cry11 toxins from *B. thuringiensis*, and the Tpp1Aa2/Tpp2Aa2 toxin from *L. sphaericus* (20–23).

Here, we employed SFX using the novel nano-focus option at the SPB/SFX instrument (24) of the European XFEL, to match beam focus to the size of natural crystals of Tpp49Aa1 isolated from recombinant *B. thuringiensis* expressing the *L. sphaericus* Tpp49Aa1 protein as parasporal inclusions. In addition to the ability of the European XFEL to deliver the highest available X-ray intensity per pulse, the unique MHz-pulse structure allowed for data collection at high repetition rate and determination of the Tpp49Aa1 structure from a high quality, high redundancy dataset. The Tpp49Aa1 structure will be important for understanding the mechanism of the Cry48Aa1/Tpp49Aa1 two-component system fully. We also model the interaction of Tpp49Aa1 with Cry48Aa1 and validate cell-based model systems to study their action *in vitro*. Given that the Cry48Aa1/Tpp49Aa1 pair exhibits toxicity against Tpp1Aa2/Tpp2Aa2-resistant *C. quinquefasciatus* larvae (7, 8), our work has implications for managing mosquito resistance and we identify an expanded range of mosquito targets for control by Cry48Aa1/Tpp49Aa1.

## 2. Results and Discussion

### 2.1. Insect Bioassays

To verify the quality of the proteins to be studied, a combination of the recombinant Cry48Aa1 and Tpp49Aa1 were bioassayed on three mosquito species that have been previously investigated. After 24 hours exposure to a high dose of the Cry48Aa1/Tpp49Aa1 pair, we observed mortality in the *C. quinquefasciatus* larvae. A high dose of Cry48Aa1/Tpp49Aa1 caused no mortality in *Ae. aegypti* or *An. gambiae*, (nor did a high dose of the individual toxin components) - confirming the target range noted in previous publications (9), and the functionality of the Tpp49Aa1 produced for this study.

We also tested the activity of the Cry48Aa1/Tpp49Aa1 pair against larvae from three mosquito species that have not been previously investigated for susceptibility: *Aedes albopictus, Anopheles stephensi*, and *Culex tarsalis*. Cry48Aa1/Tpp49Aa1 demonstrated toxicity against all three species, with LC_50_ of 111 ng/mL, 173 ng/mL, and 91 ng/mL at 48 hours, respectively (**Table 1**). This significantly enhances the known target range for Cry48Aa1/Tpp49Aa1, increasing its value in mosquito control. These mosquito species are significant vectors of human disease, with *Ae. albopictus* reported as a primary vector for emerging arboviruses (25)– such as dengue virus – and *C. tarsalis* suggested as the most important vector of human West Nile virus in the USA upper midwest region (26). Furthermore, *An. stephensi* was recently reported as a new malaria vector in Africa. *An. stephensi* is one of the few anopheline species found in central urban locations, putting 126 million Africans at risk, and prompting the World Health Organization to issue a vector alert and call for targeted control and prioritized surveillance (27).

**Table 1.** Potency of a 1:1 mixture of Cry48Aa1/Tpp49Aa1 to fourth instar mosquito larvae.

| **Mosquito Species** | **Exposure (hours)** | **LC<sub>50</sub> (ng/mL)**<br>**(95% fiducial limits)** | **LC<sub>50</sub> (ng/mL)**<br>**(95% fiducial limits)** |
| --- | --- | --- | --- |
| *Aedes albopictus* | 24 | 607 (332 – 1,110) | 4,800 (2,630 – 8,800) |
|  | 48 | 111 (68 – 183) | 1,360 (823 – 2,230) |
| *Anopheles stephensi* | 24 | 818 (450 – 1,490) | 3,830 (2,100 – 6,960) |
|  | 48 | 173 (118 – 253) | 712 (450 – 965) |
| *Culex tarsalis* | 24 | 1,890 (578 – 6,160) | 301,000 (92,100 – 982,000) |
|  | 48 | 91 (61 – 134) | 436 (295 – 646) |

### 2.2. Structure Description

Diffraction data were collected from Tpp49Aa1 nanocrystals, which had been visualized by transmission electron microscopy **(Fig. S1)**. The sample was injected in the vacuum chamber of the SPB/SFX beamline at European XFEL using 3D-printed microfluidic double flow focusing nozzles (DFFN). In total, 2,458,059 diffraction patterns were collected from which 157,582 of the strongest patterns were indexed in space group P2_1_2_1_2_1_. Phasing was performed by molecular replacement using *L. sphaericus* Tpp1Aa2/Tpp2Aa2 (PDB 5FOY and PDB 5G37) and *L. sphaericus* Tpp2Aa3 (PDB 3WA1) as starting models. Manual building and refinement from the ensembled diffraction data led to a model with R_work_/R_free_ of 0.178/0.197 at 1.62 Å resolution **(Table S1)**. The electron density map **(Fig. S2)** showed continuous density for residues 49 – 464 of the protein sequence. The first 48 N-terminal residues are not observed in the map. The structural data reveal that Tpp49Aa1 crystals are formed by the packing of dimers into the crystal lattice.

Within each Tpp49Aa1 monomer **(Fig. 1)**, two distinct domains exist: An N-terminal lectin-like head domain covering residues 49 – 214, and a C-terminal putative pore-forming domain (PFD) covering residues 215 – 464. Related proteins were identified in the Protein Data Bank using the DALI server (28) to perform a structural similarity search with Tpp49Aa1. The best matches were other pesticidal proteins belonging to the Tpp family (PF05431), sharing both the common PFD and β-trefoil domains **(Fig. 2)**: *L. sphaericus* Tpp2Aa2 (PDB 5FOY – chain B), *L. sphaericus* Tpp2Aa3 (BinB3 variant, PDB 3WA1 – chain A), *L. sphaericus* Tpp1Aa2 (PDB 5FOY – chain A), and *B. thuringiensis* Tpp35Ab1 (formerly Cry35Ab1, PDB 4JP0 – chain A), with greatest structural similarity between Tpp49Aa1 and Tpp2Aa2 **(Table S2)**.

**Figure 1.**
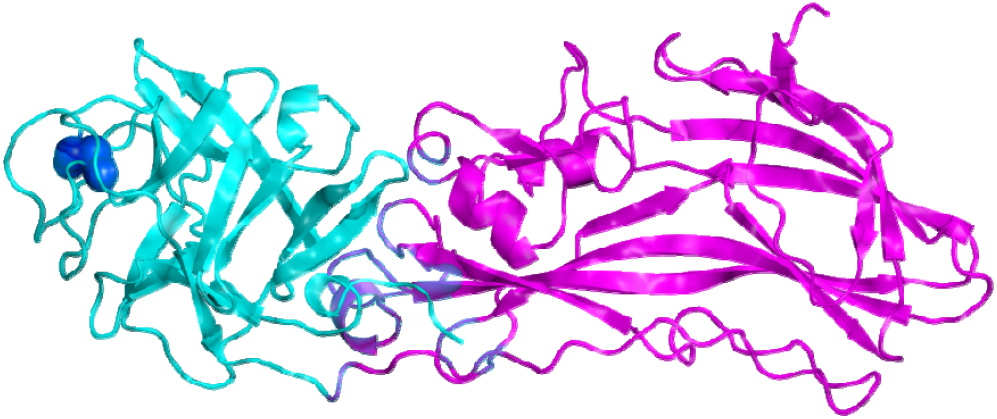
Crystal structure of Tpp49Aa1. Two distinct domains exist within the Tpp49Aa1 monomer. The N-terminal lectin-like head domain (cyan) consisting of six β-hairpins that form a β-trefoil fold containing a disulphide bond Cys91-Cys183 (shown as spheres – dark blue). The C-terminal pore forming domain (magenta) comprises the aerolysin-like domain.

**Figure 2.**
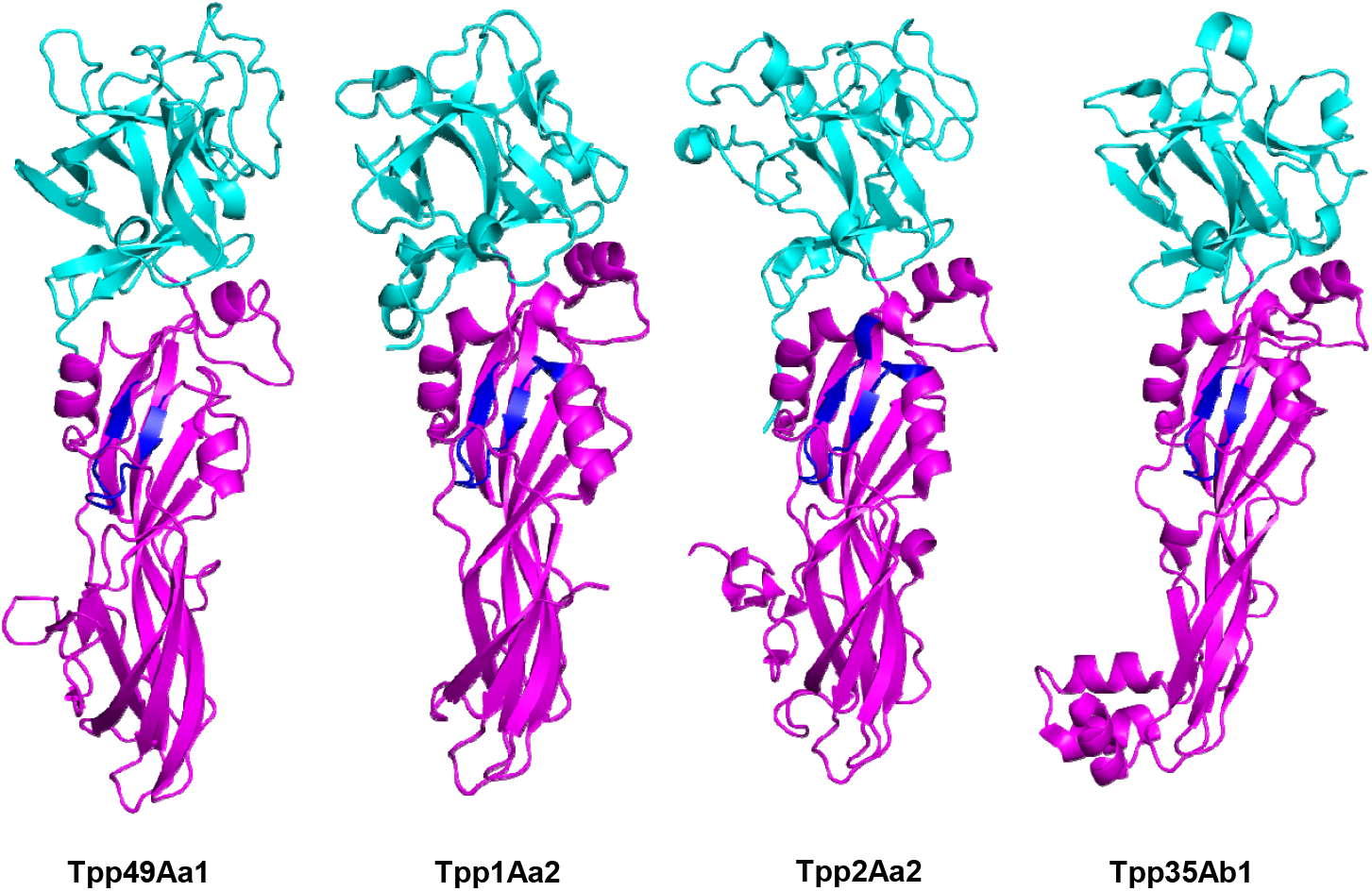
Comparative structures of β-sheet toxins. The structures of pesticidal proteins: Tpp49Aa1 (PDB 7QA1) from *L. sphaericus*, Tpp1Aa2 from *L. sphaericus* (PDB 5FOY – chain A), Tpp2Aa2 from *L. sphaericus* (PDB 5FOY – chain B), Tpp35Ab1 from *B. thuringiensis* (PDB 4JP0 – chain A). Lectin-like domains are shown in cyan and pore-forming domains are shown in magenta. Regions including a short β-hairpin with predominantly amphipathic structure (shown in dark blue) were proposed as transmembrane domains in Tpp1 (residues 256-268) and Tpp2 (residues 302-317). A similar structure is found in the structure of Tpp35 (residues 249-259) and in our Tpp49 structure (residues 322-334).

The first 48 N-terminal residues are not seen in the structure, but are known to be present in Tpp49Aa1 crystals produced in the native host *L. sphaericus* (8). To ensure that this region is also present when (as in this work) the protein is expressed in *B. thuringiensis*, N-terminal sequencing was performed for 5 cycles. The resulting sequence MXNQ(E) appears to correspond to the authentic N-terminal sequence MENQI, showing that the N-terminus is intact and, therefore, apparently disordered in the crystals. This region is known to be proteolytically removed from the protein by *C. quinquefasciatus* gut extracts (9) and the structural flexibility that might disorder the sequence in the crystals may facilitate this activation of the protein. The N-terminal domain seen in the structure comprises a β-trefoil fold consisting of six two-stranded β-hairpins which form a β-barrel with a triangular cap, as well as one disulphide bond, Cys91-Cys183. The β-hairpins in this, and other β-trefoil folds, are arranged into three subdomains commonly designated α, β, and γ, giving rise to a pseudo three-fold axis. β-trefoil folds are highly conserved in the lectin family, a class of carbohydrate-binding proteins (29). Several studies have indicated a role for carbohydrate moieties in eliciting pesticidal action of Cry proteins (30–33). Hence, it is possible that carbohydrate binding may also be important for pesticidal activity of the Cry48Aa1/Tpp49Aa1 toxin pair. Although, in Tpp2Aa2 of the Tpp1Aa2/Tpp2Aa2 complex, the Cys67-Cys161 disulphide bond, which is the equivalent of the Cys91-Cys183 disulphide bond identified in the Tpp49Aa1 structure **(Fig. S3)**, has been proposed to obstruct the putative α sugar binding module (21), elimination of the Cys residues in either Tpp2Aa3 (34) or in Tpp49Aa1 (15), eliminates toxicity.

The C-terminal domain of Tpp49Aa1 comprises a β sheet-rich topology characteristic of the aerolysin family of pore-forming toxins (35, 36). Indeed, Lacomel *et al*. speculated that the Tpp family may represent a subclass of the larger aerolysin, ETX/MTX-2 superfamily of β-pore forming proteins (37). Within this superfamily, the aerolysin toxin is considered the model member and its mechanism of pore formation has been well characterised. Aerolysin, which is secreted as the inactive proaerolysin homodimer, is first activated by proteolytic cleavage of the C-terminal propeptide, allowing dissociation to a monomer (35, 38). The C-terminal PFD interacts with target glycosylphosphatidylinositol anchored receptors, whilst the N-terminal receptor binding domain interacts with target glycans (39). Oligomerization of seven monomers, via interaction of the PFDs, leads to formation of a pre-pore structure constituting an amphipathic β-barrel, which inserts into the membrane, leading to cell death by osmotic lysis (38). The structural homology seen between Tpp49Aa1 and the aerolysin family may suggest that Tpp49Aa1 is able to form pores by a similar mechanism. In an analysis of the Tpp proteins and the wider aerolysin family (37), a region including a short β-hairpin with predominantly amphipathic structure, tucked under a loop within the PFD, was proposed as the transmembrane domain in Tpp1 (residues 256-268) and Tpp2 (residues 302-317) **(Fig. 2)**. A similar structure is found in the structure of Tpp35Ab1 (residues 249-259) and in our Tpp49Aa1 structure (residues 322-334) **(Fig. 2)**. We speculate that, as in an aerolysin-like mechanism, this region may unfold to form the beta barrel pore in the target cell membrane. However, it also cannot be ruled out that pore formation by Tpp49Aa1 may occur via unique mechanisms and the necessity for Cry48Aa1 in the action of Tpp49Aa1 raises further questions regarding the formation of pores by this protein pair. Additional structural studies are required to investigate the pore-formation process further.

The final model of Tpp49Aa1 revealed the presence of a homodimer forming an “X” structure with a large intermolecular interface **(Fig. 3)**, similar to that described for natural heterodimeric crystals of Tpp1Aa2/Tpp2Aa2 **(Fig. S3)** (21). Superposition of the Tpp49Aa1 monomers shows the two copies to be almost identical, with an all-atom RMSD of 0.681 Å **(Fig. S4)**. The interface between the two monomers, which involves 41 residues from monomer A and 42 residues from monomer B, forming 16 hydrogen bonds **(Table S3, Fig. 3)**, exhibits an area of 1329.1 Å^2^ and the predicted binding energy is -11.1 kcal mol^-1^ **(Table S3)**. In the Tpp1Aa2/Tpp2Aa2 heterodimer **(Fig. S3)**, the interface between the monomers involves 49 residues from Tpp1Aa2 and 63 residues from Tpp2Aa2, making 19 hydrogen bonds and 2 salt bridges. Interface analysis by PDBePISA estimates the area at 1833.1 Å^2^ and the binding energy at -22.5 kcal mol^-1^, indicating a more stable complex for Tpp1Aa2/Tpp2Aa2 heterodimers than for Tpp49Aa1 homodimers. Indeed, whilst interaction of the solubilised and activated forms of Tpp1Aa2/Tpp2Aa2 is required for full toxicity in the mosquito larvae, interaction of Tpp49Aa1 may only occur in the crystal to support packing and stability. These different interactions highlight an interesting feature of the Tpp family, in which some members (Tpp36, Tpp78, Tpp80) can exert pesticidal activity alone, while Tpp1/Tpp2 requires interaction of two distinct members of the Tpp group, Tpp35 requires interaction with Gpp34, and Tpp49Aa1requires interaction with Cry48Aa1, a member of a distinct (Cry) structural family.

**Figure 3.**
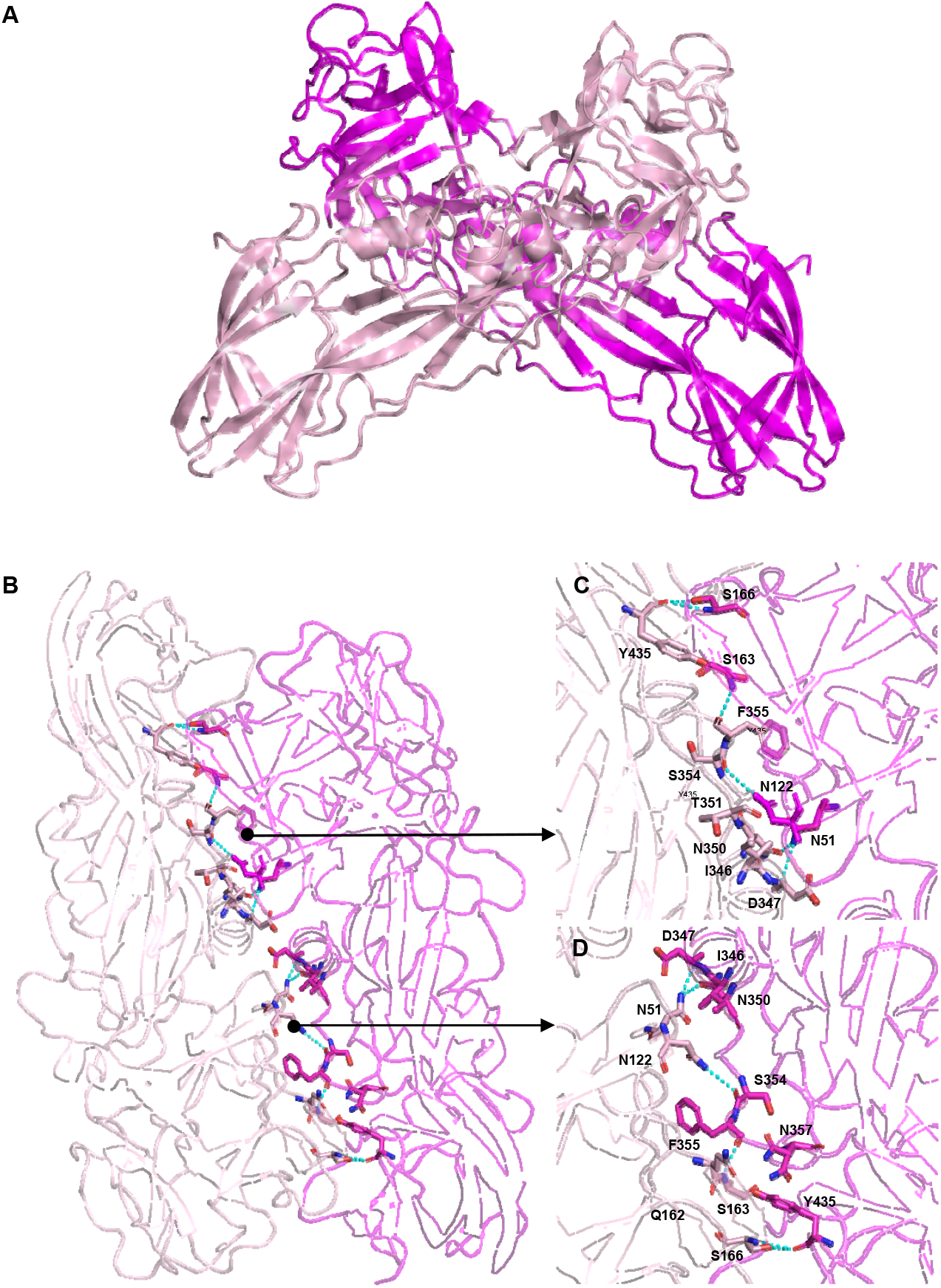
Structure of Tpp49Aa1 homodimer. **(A)** Tpp49Aa1 homodimer (monomer A – magenta, monomer B – light pink) forming an “X” structure with a large intermolecular interface. **(B)** Tpp49Aa1 dimer interface. Residues involved in hydrogen bonds identified by PDBePISA are shown as sticks (carbon – magenta / light pink, oxygen – red, nitrogen – blue). Polar interactions within a 3.6 Å cut-off are highlighted by cyan dashes. **(C, D)** Detailed view of interfacial residues involved in hydrogen bonding.

To investigate whether the Tpp49Aa1 dimeric form is maintained in solution, solubilised Tpp49Aa1 was assessed using SEC (Supplementary methods 1). Tpp49Aa1 is present as two major bands that run at, and just above, the 48 kDa marker on an SDS-PAGE gel (**Fig. S5.A**). Solubilised Tpp49Aa1 crystals have previously been shown to produce similar size bands (8), and these may be products that are produced during analysis by bacterial proteinases contaminating the crystal preparation. SEC showed a main peak at 51 kDa, but with a shoulder peak before at 121 kDa indicating that it is predominantly a monomer but that some dimers may persist (**Fig. S5.B**). In further analysis, static light scattering and refractive index measurements were used to analyse the molecular weight of the protein-containing fractions (Supplementary methods 2). Calibrated to BSA, this indicated Tpp49Aa1 is present in solution at a single MW corresponding to 52.1 kDa, approximately the weight of the monomer (**Fig. S5.C**). It is possible that after initial SEC and concentration of protein-containing peaks, the equilibrium between monomeric form and dimeric form is altered. Within the natural crystals, the N-terminal region of Tpp49Aa1 (residues Asn49 – Ser148) that has been shown to interact with Cry48Aa1 (12), is partially buried within the large intermolecular interface **(Fig. S6)**. Dissociation of the Tpp49Aa1 dimer would, therefore, expose this region for interaction with Cry48Aa1. Both methods discussed above indicate that Tpp49 is predominantly present as a monomer in solution.

### 2.3. Alkaline-triggered release of Tpp49Aa1

Another important feature that Tpp49Aa1 shares with a range of other pesticidal proteins in the taxonomic class Bacilli, is its propensity to form natural crystals when expressed at high level (in its source organism, *L. sphaericus*, and in this study, in recombinant form in *B. thuringiensis*). Tetreau *et al*, have recently reviewed factors within proteins that may contribute to the ability of bacteria to sequester them into natural crystals (40). Natural crystals of Cry3Aa (PDB 4QX0) (20) were found to have a high solvent content: 60.4% by Matthews’ analysis compared to a mean of ∼47% in a previous analysis of *in vitro* grown crystals of proteins in the PDB (41). Tpp49Aa1 has an estimated 49.6% solvent content - closer to the mean of *in vitro* grown crystals - but, like natural crystals of Cry3Aa, the Tpp49Aa1 crystals are permeated by wide solvent accessible channels **(Fig. S7)**. We can speculate that the solvent channels within the crystals may facilitate dissolution in the appropriate environment.

The ability of natural crystals to remain stable in the open environment but to be dissolved by insect gut pH changes is thought to rely on a range of factors, including intermolecular salt bridges that may be pH labile, intermolecular hydrogen bonds that may be deprotonated upon pH elevation leading to electrostatic repulsion, and other gut solutes that may affect this solubilisation. PDBePISA analysis shows that the packing of dimers into crystals introduces eight possible interfaces between neighbouring dimers **(Table S3)**. A network of hydrogen bonds and salt bridges is also observed. Given that Tpp49Aa1 crystals are readily solubilised in the alkaline environment of the mosquito larval gut, we next sought to investigate the effect of extreme pH on Tpp49Aa1 crystals. To do so, pH 3 or pH 11 buffers were injected into the vacuum chamber of the SPB/SFX beamline, allowing mixing with the natural Tpp49Aa1 crystals prior to data collection. In total, 466,741 / 707,992 diffraction patterns were collected from which 279,362 / 426,506 could be indexed in space group P2_1_2_1_2_1_, (a∼79.89 / 80.12; b∼82.84 / 83.22; c∼157.88 / 156.49 Å; α=β=□=90) for the pH 3 / 11 datasets, respectively **(Table S1)**. Phasing was performed by molecular replacement using the native (pH 7) Tpp49Aa1 structure as the starting model. Manual building and refinement from the ensembled diffraction data led to a model with R_work_/R_free_ of 0.189/0.213 at 1.78 Å resolution for data collected at pH 3, whilst a model with R_work_/R_free_ of 0.177/0.199 at 1.75 Å resolution was obtained from data collected at pH 11 **(Table S1)**. In both models, the electron density map **(Fig. S2)** showed continuous density for residues 49 – 464 of the protein sequence. As in the map obtained for the pH 7 dataset, the first 48 N-terminal residues are not observed in the pH 3 / 11 maps.

The pH 7 and pH 11 structures align closely, displaying an all-atom RMSD of 0.487 Å. As in the Tpp1Aa2/Tpp2Aa2 structure (21), an increase in pH from 7 to 11 is estimated to alter the net charge of Tpp49Aa1 from -6.518 to -47.822 e, suggesting that alkalinity will destabilize the crystal by negative electrostatic repulsion. Within the Tpp49Aa1 monomer, the largest conformational changes occur within the surface-exposed loops of the Tpp49Aa1 monomer, suggesting that these regions are affected by pH. Alkalinity also perturbs the packing of Tpp49Aa1 dimers into crystals, identified by analysis of crystal contacts and interfacial interactions using PDBePISA **(Tables S3, S4, S5)**. Overall, the binding energy of all eight crystal contacts (excluding the dimer interface) increases by 1.8 kcal mol^-1^, indicating a weaker affinity. Despite this, the binding energy of the dimer interface decreases (pH 7 = -11.1, pH 11 = -11.9 kcal mol^-1^), indicating an increase in affinity. This may suggest that, during dissolution, contacts between individual dimers dissociate prior to the dimers themselves. Subsequent release of the Tpp49Aa1 monomers will expose the regions required for activation and interaction with the target membrane and partner protein, Cry48Aa1.

Alkalinity also induces the loss or formation of 18 interfacial interactions at both the dimer interface and crystal contacts **(Table S5)**. The most notable changes include loss of hydrogen bonds Asn51[ND2]-Asn350[OD1] and Asn51[ND2]-Asp347[O] of both monomers at the dimer interface. Given the proximity of these interactions to the cleavage site (Phe48) of the N-terminal pro-peptide, we hypothesize that loss of these hydrogen bonds would increase accessibility of the pro-peptide to proteases. In addition, elevated pH leads to loss of a salt bridge, Asp429(OD1)- Arg267(NH1), at a crystal contact outside of the dimer interface, supporting the idea that crystal stability is thought to rely, in part, on intermolecular salt bridges. Other changes in the interactions have been listed in **Table S6A**. Similar changes were seen following a decrease in pH from 7 to 3 **(Table S6B)**, although this is not physiologically relevant to pH values found in the target insect guts (although this low pH is capable of solubilising crystals). Briefly, mixing at pH 3 results in a larger increase in overall binding energy at the eight crystal contacts (5.0 kcal mol^-1^) in comparison to mixing at pH 11 (1.8 kcal mol^-1^), suggesting that crystals may dissolve more readily at low pH. These studies shed light on the early events leading up to the dissolution of natural Tpp49Aa1 crystals.

### 2.4. Modelling the Cry48Aa1-Tpp49Aa1 interaction

Mosquito toxicity bioassays have revealed that both Cry48Aa1 and Tpp49Aa1 in combination are required for activity against *C. quinquefasciatus*, with optimal activity arising from a ratio of ∼1:1 (8, 9). In addition, dot blot assays have revealed the ability of Cry48Aa1 and Tpp49Aa1 to form a complex (12). Given these studies, it is interesting to speculate how Tpp49Aa1 and Cry48Aa1 might interact.

The full-length Cry48Aa1 structure was predicted using the AlphaFold2 package (42) and the core toxin domains (residues 53 – 659) were extracted for docking. The suitability of the Alphafold2 package to predict the structures of pesticidal proteins was assessed by predicting the Tpp49Aa1 structure elucidated here and showed an all-atom RMSD of 1.982 Å of the predicted and actual structures **(Fig. S8)**. To model the Cry48Aa1-Tpp49Aa1 complex, we used the ClusPro and RosettaDock programs. Briefly, ClusPro was employed to perform a global docking search, from which the central models of the top 5 clusters were carried forward for refinement by performing local docking searches using RosettaDock. From each of the 5 local docking searches, 1,000 models were produced. The thermodynamic hypothesis states that models with the lowest energy scores are most likely to represent that of the native structure (43). Therefore, within each of these searches, output models were ranked according to their total energy scores and the 5 lowest scoring models were then identified. Models are referred to according to their ClusPro cluster (c1 – c5) followed by their Rosetta energy rank (r1 – r5, with r1 being the lowest energy score and r5 being the highest energy score) **(Table S7)**. To analyse the interfacial interactions, PDBePISA was utilized. Interface area ranged from 478.8 to 1101.1 Å^2^ and all models were found to contain several interfacial hydrogen bonds, with the interfaces of some models also containing salt bridges **(Table S7)**.

In addition to analysing the interfacial interactions of modelled Cry48Aa1-Tpp49Aa1 complexes, structural stability was assessed. To do so, 100 ns molecular dynamics (MD) simulations were performed (Supplementary methods 3) and the RMSD of the position of backbone atoms was calculated as a function of time. MD simulations enable the physical movement of atoms and molecules to be analysed. Low RMSD values indicate stable structures which remain close to their starting positions, whilst high RMSD values indicate unstable structures which deviate away from their starting positions. This approach relates to the free energy landscape model which assumes that the correctly docked conformation exists at the bottom of the funnel within the free energy landscape (44). Similar approaches have been used previously and yielded models consistent with experimental structures (45–47). Here, several models originating from ClusPro clusters 1, 3 and 4 were found to exhibit RMSD values that remained relatively constant throughout the 100 ns MD simulations **(Fig. S9.A, S9.C, S9.D)**. Of these models, only clusters 3 and 4 exhibited binding consistent with previous work indicating that the N-terminal region of Tpp49Aa1 (Asn49-Ser148) is responsible for binding Cry48Aa1 (12). Of these two clusters, models c3r3 **(Fig. 4.A)** and c4r3 **(Fig. 4.B)** were found to exhibit the lowest RMSD values **(Fig. S9.C, S9.D)**. In model c3r3, Cry48Aa1 interacts via domain II (β-sheet domain) and domain III (antiparallel β-sandwich domain) with both the N- and C-terminal domains of Tpp49Aa1 - but domain I is not involved. This contrasts with model c4r3, where only the Tpp49Aa1 N-terminal domain interacts, forming contacts with Cry48Aa1 mainly in domain I (minor interactions with domains II and III). The c4r3 model is also consistent with the findings of Guo et al. 2020 (48), showing that Tpp49Aa1 is able to interact with fragments of Cry48Aa1 containing domain I (residues 53-239) and fragments spanning domains II and III. The binding energy (calculated using PISA) of model c3r3 is -7.5 kcal mol^-1^ (with -5.9 kcal mol^-1^ from interaction with the Tpp49Aa1 C-terminal domain and only -1.6 kcal mol^-1^ from interaction with the Tpp49Aa1 N-terminal domain). In contrast, the binding energy of model c4r3 - from binding via just the Tpp49Aa1 N-terminal domain is -9.1 kcal mol^-1^, consistent with the fact that the N-terminal residues Asn49-Ser148 alone are sufficient for Cry48Aa1 interaction (12). Model c4r3 is further supported by a study demonstrating that mutations at Cys91 and Cys183, present at the Cry48- Tpp49 interface in this model and shown to form a disulphide bridge in the Tpp49Aa1 structure, lead to weaker Cry48Aa1 interaction (15). As a result, we propose model c4r3 as the most favoured model of the interaction (**Fig. 4.B**).

**Figure 4.**
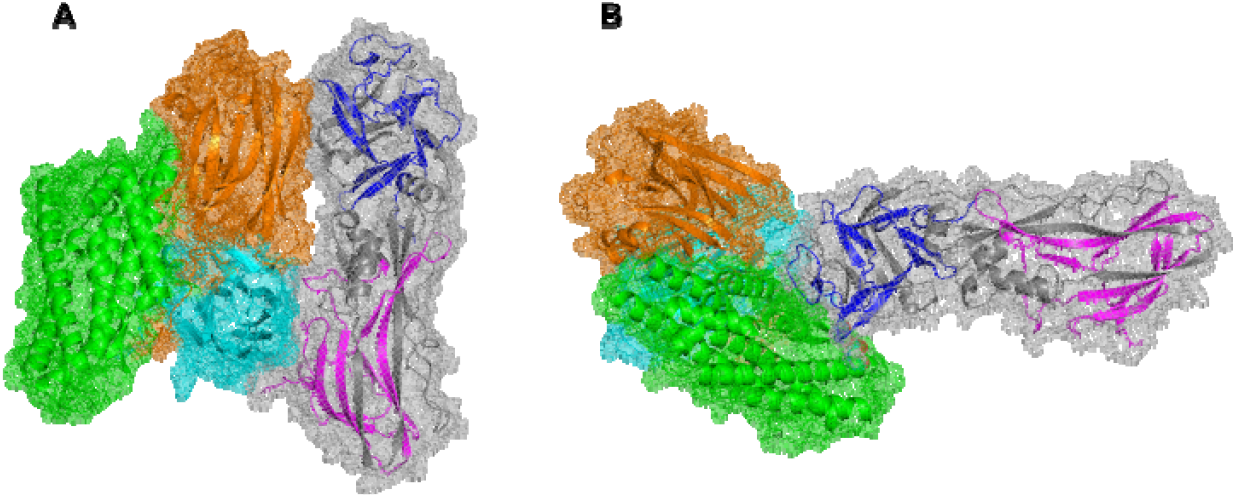
Cry48Aa1-Tpp49Aa1 models. The top 2 predicted dimer structures are illustrated. Cry48Aa1 contains three domains shown in green (domain I), orange (domain II) and cyan (domain III). Tpp49Aa1 is coloured grey, with regions experimentally shown to interact with Cry48Aa1 (blue) and the cell membrane (magenta) also highlighted. **(A)** Model c3r3 and **(B)** model c4r3.

### 2.5. Cellular models of Cry48Aa1 and Tpp49Aa1

The cellular toxicity of Cry48Aa1 and Tpp49Aa1 was investigated in two cell lines derived from the known target species *C. quinquefasciatus* (MRA-918 and Hsu), and a cell line from the newly identified target - *C. tarsalis* (Ct). To investigate cytotoxicity, resazurin was used as an indicator of metabolic activity and cellular viability (49). Cry48Aa1-Tpp49Aa1 treatment reduced cellular metabolism in all three cell lines, confirming a functional effect of the proteins generated in this study (**Fig. 5**).

**Figure 5.**
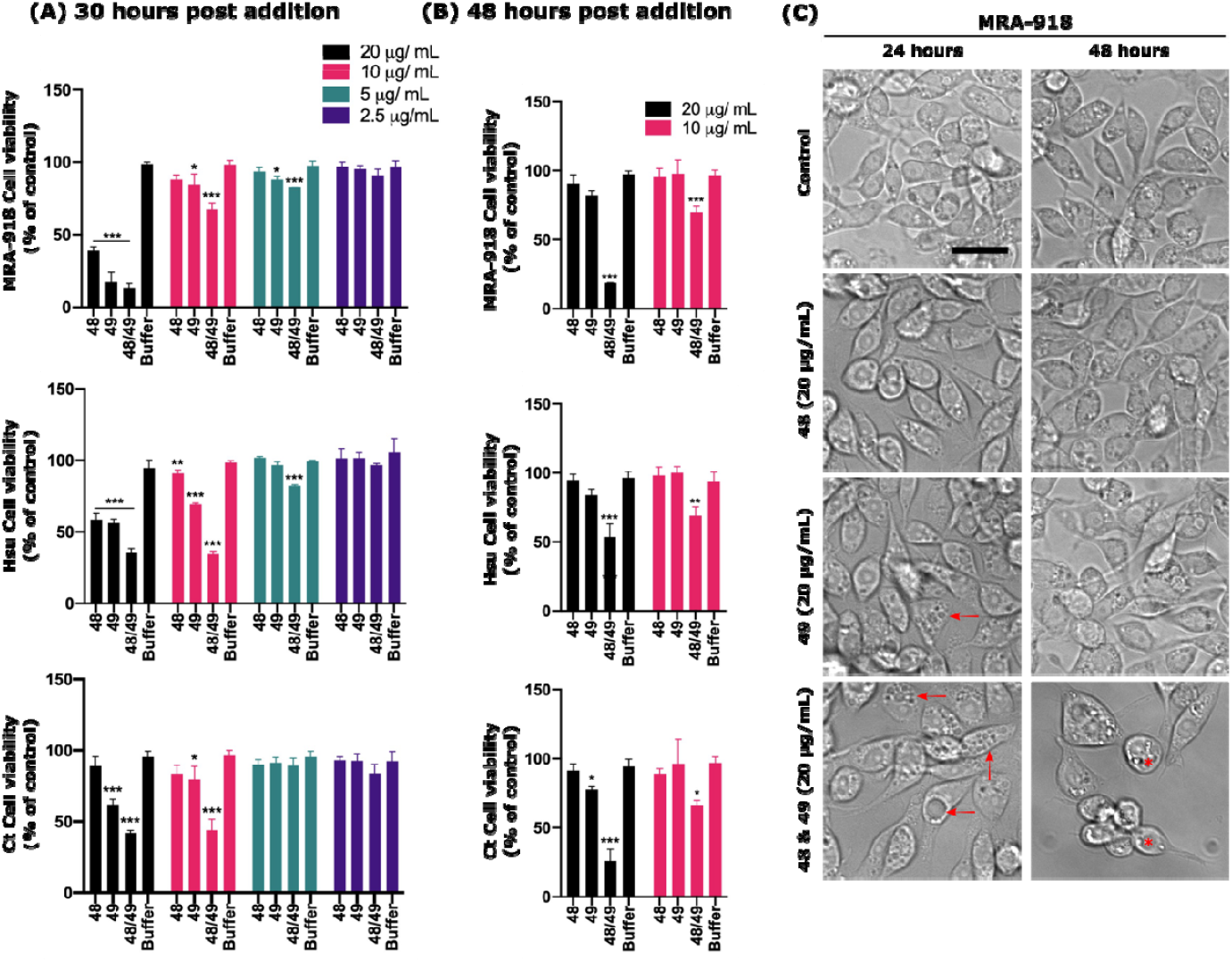
Cry48Aa1 and Tpp49Aa1 reduce cell viability in *C. quinquefasciatus* and *C. tarsalis* cell lines. To assess the use of various insect cell lines for modelling Cry48Aa1 and Tpp49Aa1 toxicity, resazurin was used to quantify the effect of these proteins on cell viability. *C. quinquefasciatus* (MRA-918, Hsu) and *C. tarsalis* (Ct) cells were treated with a range of concentrations of either Cry48Aa1 (“48”), Tpp49Aa1 (“49”), or equimolar amounts in combination (48/49). Resazurin was added to the cells 24 h **(A)** and 48 h **(B)** after the pesticidal proteins. **(A)** 24-hours post exposure, reduced cell viability (% of control) was observed in all cell lines, with either 48/49 in combination or alone. **(B)** Twenty-four hours post 48/49 exposure, reduced cell viability is still seen with both pesticidal proteins (alone and in combination), however, the decreased viability between the single protein treatments is reduced compared to the combination, indicating recovery. All data are presented as the mean ± SD and all statistical analysis was performed using a one-way ANOVA, n = 3 (***p < 0.001, **p < 0.01, * p < 0.05). **(C)** Brightfield images of MRA-918 cells at 24 h, post pesticidal protein addition, show a slightly rounded cellular morphology with either 48 or 49, alongside the presence of some small vacuoles in 49 alone (red arrows). Larger vacuoles are apparent in 48/49 combination (red arrows). At 48 h post 48/49 addition, cells have rounded and detached from the plate (red asterix). Representative scale bar = 10 μm.

In cells from the previously known target species, *C. quinquefasciatus*, higher concentrations (20, 10, & 5 μg/mL) of the Cry48Aa1-Tpp49Aa1 combination reduced metabolic activity after 24 h in both cell lines (**Fig. 5.A**). By 48 h, cells subjected to the higher concentrations (20 & 10 μg/mL) still showed reduced viability (**Fig. 5.B**). When added as individual components, both Cry48Aa1 and Tpp49Aa1 caused reduced viability, as quantified at 24 h (**Fig. 5.A**). However, by 48 h, cells treated with individual proteins recovered, with no difference in viability compared to controls. This suggests both components are required for long-term cytotoxicity and cell death – although whether this is through formation of a complex remains to be determined. Similar concentration-dependent effects on cell viability were noted in the *C. tarsalis*-derived cell line (Ct), with both toxin components required for a significant, sustained decrease in viability at 48 hours post exposure, with individual toxin components having a small effect on cell viability at 24 hours (**Fig. 5.A**), that recovered at 48 hours post exposure (**Fig. 5.B**). Whether these lesser, recoverable, effects of a single component are mediated through the same cellular pathways as the Cry48Aa1-Tpp49Aa1 pair is yet to be determined.

Light microscopy of the MRA-918 cell line shows the Cry48Aa1/Tpp49Aa1 combination - and to a lesser degree Tpp49Aa1 alone – can induce vacuolisation at 24 hours post exposure (**Fig. 5.C**). This phenotype appears to recover in the Tpp49Aa1 only treated cells at 48 hours post exposure, whereas in the Cry48Aa1-Tpp49Aa1 treatment we observed a rounded morphology, retraction of cellular processes and significant detachment from the plate, indicative of cell death. In agreement with our viability assays, this indicates both components are required for full toxicity, yet Cry48Aa1 or Tpp49Aa1 alone may have a small, temporary effect on cellular health. Cytoplasmic and mitochondrial vacuolisation has been reported in previous studies administering Cry48Aa1/Tpp49Aa1, in combination, to midgut epithelial cells isolated from *C. quinquefasciatus* larvae (14). Similar phenotypes are also reported with Tpp49Aa-related Toxin_10 family members Tpp1Aa1/Tpp2Aa2 in both larvae (50, 51) and cells expressing the relevant Cpm1 receptor (52).

All three *Culex* cell lines may be useful models for investigating the cellular pathways through which Cry48Aa1-Tpp49Aa1 toxicity is elicited. No effects on the metabolic activity or cellular morphology were apparent from addition to Sf9 cells, either in combination or as individual proteins, consistent with the fact that Cry48Aa1-Tpp49Aa1 is reported non-toxic to the source insect - *Spodoptera frugiperda* (**Fig. S10**). Interestingly, the concentration of Cry48Aa1-Tpp49Aa1 required to elicit cell death in *Culex* cell lines is approx. 1000-fold higher than that previously reported to exert death in *C. quinquefasciatus* larvae. This may be due to the tissue origin of the cells – Hsu is an ovary derived cell line and MRA-918 are from neonate larvae. We know the site of action for these toxins is the mosquito midgut, and the expression level of the relevant receptor(s) in the cell lines is yet to be determined. The toxin pair clearly displays some insect cell line selectivity and the necessity of both components for full cytocidal activity is clear – yet further work is required to know if the mechanisms of cell death are the same *in vitro* as *in vivo*.

## 3. Conclusions

We have used megahertz SFX at the European XFEL to determine the Tpp49Aa1 structure to a final resolution of 1.62 Å. Complementary experiments conducted at varied pH also enabled investigations of the early structural events leading up to the dissolution of Tpp49Aa1 crystals, shedding light on the mechanisms of crystal dissolution. It has been shown that the nano-focus option at the SPB/SFX instrument, in combination with megahertz repetition rates, can be used for rapid and high-quality data collection from natively grown nanocrystals and this paves the way for further investigations on structure and dynamics of bacterial insecticides. In addition, we have expanded the target range of the Cry48Aa1/Tpp49Aa1 protein pair to include *Ae. albopictus, An. stephensi* and *C. tarsalis* and identified MRA-918, Hsu, and Ct cell lines as useful models for investigating the cellular pathways by which toxicity may be elicited. In the future, structural investigations of Cry48Aa1, both in the crystalline protoxin and activated forms, and the Cry48Aa1-Tpp49Aa1 complex, will shed further light on the mode of action of these toxins and aid the design of optimized and safe insecticides to combat mosquitoes as vectors of a range of neglected tropical diseases.

## 4. Materials and Methods

### 4.1. Purification of Tpp49Aa1 Protein Crystals

The *B. thuringiensis* recombinant strain 4Q7::pHTP49, encoding the *tpp49Aa1* gene (accession number AJ841948) (8) was grown in 400 mL Embrapa medium (53) containing 5 μg/Ml erythromycin at 30°C with shaking (200 rpm) until sporulation reached > 90%, as judged by phase contrast microscopy. Sporulated cultures were harvested and the natural crystal proteins were isolated using stepped sucrose gradients as previously described (8). Crystal protein was run on SDS-PAGE and blotted onto PVDF membrane for N-terminal sequencing by Alta Bioscience Ltd (Redditch, UK).

For use in bioassays, Cry48Aa1 crystals were prepared from recombinant Bt strain 4Q7::pSTABP135 (8) using the method described above.

### 4.2. Bioassays

Bioassays were carried out against a range of insects (*C. quinquefasciatus, Ae. aegypti* and *An. gambiae, Ae. albopictus, An. stephensi, and C. tarsalis*) and insect cell lines including *C. quinquefasciatus* (Hsu, and MRA-918) *C. tarsalis* (Ct), and *S. frugiperda* (Sf9) (54–56). It should be noted that the MRA-918 line (also known as 4A3A) was originally reported as an *An. gambiae* line but the cells have been independently verified by a number of laboratories, including our own (amplifying and sequencing mitochondrial large ribosomal subunit, mitochondrial cytochrome C oxidase and maltase genes), to be *C. quinquefasciatus*.

To check our toxin preparations for activity, a high dose of Tpp49Aa1/Cry48Aa1 was given either together or separately to *C. quinquefasciatus, Ae. aegypti* and *An. gambiae*. Toxins were added to 1 mL of water containing 5 third instar larvae that were checked for mortality 24 hours later. To investigate new potential targets for Tpp49Aa1/Cry48Aa1, *Ae. albopictus, An. stephensi* and *C. tarsalis* larvae were bioassayed using 350 ml cups, each containing 100 ml of distilled water with 10 fourth instar larvae. A range of concentrations of 1:1 ratio w/w Cry48:Tpp49 were tested, one concentration per cup, with three replicates per concentration. Mortality was determined at 24 and 48 hours. LC_50_ and LC_95_ values were determined using Finney’s probit analysis.

Insect cell lines were maintained at 27°C in an appropriate growth medium which was changed every 4 days: Hsu, MRA-918, Ct (Schneider’s Insect Medium supplemented with 10% FBS) and Sf9 (Grace’s Insect Medium supplemented with 10% FBS). For cellular bioassays, cells were plated at 10,000 cells per well of a 96-well plate in 150 μL of medium and left to reach ∼80% confluency. Tpp49Aa1/Cry48Aa1 proteins were solubilised in 50 mM Na_2_CO_3_ pH 10.5 + 0.05% β-mercaptoethanol. Solubilised toxin was treated with immobilized TPCK Trypsin (Thermo Scientific, 20230) overnight at 37°C, followed by separation from the protein sample by centrifugation, and added either separately, or at a 1:1 molar ratio, in a range of concentrations (20, 10, 5, 2.5 μg/mL total protein) with the equivalent amount of solubilisation buffer added into control wells (at no more than 5% of total well volume). As a measure of cell viability, resazurin (10 μg/mL) was added into the cell medium (10% v/v) at 24 h or 48 h post Cry48Aa1 and Tpp49Aa1 addition. Fluorescence was quantified using a Molecular Devices Spectramax Gemini EM plate reader (λex = 445 nm: λem = 585 nm), 6 hours post resazurin addition. Statistical analyses were performed using GraphPad Prism for Mac OS (Ver 8.2.0), using one-way ANOVAs followed by Dunnett’s multiple comparisons test to compare individual treatment groups back to the control. Data are presented as mean ± standard deviation. For imaging, MRA-918 cells were seeded in 8-well Ibidi chamber slides (Thistle, IB-80826). Brightfield images were acquired with a Zeiss AX10 inverted microscope with Axiocam Mrm camera and Axiovision 4.5.2 software (Zeiss, Cambridge, UK).

### 4.3. Transmission Electron Microscopy

Purified crystal batches were characterized using transmission electron microscopy (JEM 2100-Plus, JEOL) in the XBI lab of European XFEL (57). Holey carbon copper grids (Quantifoil R1.2/1.3) were glow discharged (GloQube Plus, Quorum Technologies) freshly before use. Crystal slurry (2 μL) was applied onto the grid and incubated for 30 s and blotted using filter paper (Whatman #1). Samples were negatively stained by placing the grid on a droplet containing 2 % (w/v) uranyl acetate and blotted immediately. Grids were placed on a second uranyl acetate droplet and incubated for 20 seconds before blotting again and left for drying on filter paper. Samples were imaged with the TEM at 200 kV acceleration voltage using an Emsis Xarosa camera in imaging and selected area electron diffraction mode.

### 4.4. Structure Determination

Crystals were washed with ddH_2_O and filtered through a cascade of nylon mesh filters (Sysmex Celltrics), ranging from 100 μm down to 5 μm mesh size. The crystal suspension was centrifuged at 200 x *g* for one minute and the supernatant – containing the Tpp49Aa1 nanocrystals – was subjected to another cascade of filtration and washing before transfer to the high-pressure sample reservoirs for injection into the XFEL beam. For pH studies, 0.1 M sodium citrate (pH 3.0) and 0.1 M sodium carbonate (pH 11.0) buffers were used. Buffers were transferred to the high-pressure sample reservoirs for injection and mixed with the Tpp49Aa1 crystals 1 m upstream of the XFEL beam, approximately 1 minute before probing with X-rays. Megahertz serial femtosecond crystallography (58, 59) diffraction data were collected at the SPB/SFX (24), instrument of the European XFEL facility, Hamburg, Germany, using fast liquid-jet based injection (60) with 3D-printed (61) DFFN (62). With this set-up, 202 images per X-ray pulse train (with 10 trains/s repetition rate) were recorded with the AGIPD Detector at an intra-train pulse rate of 0.564 MHz. A photon energy of 9.3 keV with an average of 4 mJ / pulse was delivered to the instrument, focused to a spot size of about 300 nm diameter using the Nanoscale-focusing KB optics (63), providing about 6×10^12^ photons/μm^2^/pulse at the sample. The online crystal diffraction ‘hit-rate’ was monitored using OnDA program (64) with raw data processing largely following the method described by Wiedorn et al. (58). Hit finding was performed using the program Cheetah (65) with careful optimization of the peak search parameters (--peaks=peakfinder8 --min-snr=5 -- max-res=800 --threshold=500 --min-pix-count=1 --max-pix-count=50 --min-peaks=10 --local-bg-radius=3) and masking of bad pixels. Meaningful diffraction patterns were then indexed using CrystFEL (66, 67) version 0.10.1 (--int-radius=2,4,6, --multi) using the indexing method XGandalf (68). For the detector geometry optimization the program Geoptimiser was used (69). Merging and scaling of the integrated reflection intensities was performed using the Partialator program from the same CrystFEL package (--model=xsphere --min-res=3 --push-res=1.0 for pH 3 and pH 11, --model=unity --min-res=2.5 --push-res=inf for the native (pH 7) dataset). The solvent content and number of molecules in the asymmetric unit was estimated using Matthews’ analysis, available through the MATTPROB web server (41). The phasing pipeline MRage in Phenix (70, 71) was used for initial phasing, using the sequence information and a component stoichiometry of two as input. MRage used models of *L. sphaericus* Tpp1Aa2/Tpp2Aa2 (PDB 5FOY and PDB 5G37) and *L. sphaericus* Tpp2Aa3 (BinB variant, PDB 3WA1) as templates for molecular replacement. The initial model was optimized using phenix.phase_and_build, followed by another round of Phaser and automatic model building using phenix.autobuild (72). The resulting model and maps were inspected manually using coot (73), followed by iterative refinement and model building cycles using phenix.refine (74) and coot respectively. In the initial molecular replacement, two non-crystallographic symmetry (NCS)-related domains A1 and A2 were switched (A1B2, A2B1), this error was corrected in coot and the residues linking A and B were built manually. This model was subjected to another cycle of phenix.autobuild, followed by iterative refinement and model building cycles using phenix.refine and coot respectively. Final refinement was carried out using Refmac5 (75) in the CCP4i2 package (76) keeping the R_free_-Flags generated in Phenix.

### 4.5. Modelling the Cry48Aa1-Tpp49Aa1 interaction

#### 4.5.1. Structure preparation

The full-length Cry48Aa1 structure was predicted using the AlphaFold2 package (42), as installed at DESY and available through the DESY Maxwell-cluster. For docking, the core Cry48Aa1 toxin domains after proteolytic activation (residues 53 – 659, all of which display good pLDDT confidence scores – **Fig. S8.C**) were extracted from the predicted structure. Similarly, the structure of Tpp49Aa1 was obtained by extracting the core toxin domains after proteolytic activation (residues 49 – 464) from chain B of an earlier crystal structure (PDB 7QA1) elucidated at pH 7 as part of this work. Given that dot blot interaction of the Cry48Aa1 and Tpp49Aa1 proteins was performed in PBS (12), and toxicity was shown against mosquito cell lines maintained at pH 7 – 7.4, docking studies were performed at neutral pH, with the protonation states of titratable residues reflecting this.

#### 4.5.2. Molecular docking

A naïve molecular docking approach with no presumed Cry48Aa1-Tpp49Aa1 interface was used. First, a global docking search was performed using the ClusPro web server (77). Briefly, ClusPro employs a fast Fourier transform (FFT)-based algorithm to perform rigid body docking. The 1,000 lowest energy models are clustered according to their root-mean-square deviation (RMSD) and refined via energy minimisation. ClusPro outputs the central model from each cluster, with the largest clusters ranking highest. Here, some 30 clusters were identified, with the central models from the 5 largest clusters carried forward for local docking refinement.

Local docking refinement was performed using the RosettaDock algorithm (78). Briefly, RosettaDock employs a Monte Carlo based algorithm to perform rigid body docking and side chain optimisation from a user perturbed starting position. Models are scored using the ref2015 score function (79), which is composed of several weighted terms, enabling the total energy to be calculated in Rosetta Energy Units (REU). To ensure side chains were in their lowest energy conformations, models were prepacked using the *docking_prepack_protocol.macosclangrelease* executable. Subsequently, docking was performed using the *docking_protocol.macosclangrelease* executable. For each docking search, 1,000 models were generated and the 5 models with the lowest energy scores (a total of 25 models across all 5 RosettaDock searches) were carried forward for further structural analysis.

#### 4.5.3. Molecular dynamics

Structural stability of modelled Cry48Aa1-Tpp49Aa1 complexes was assessed by performing molecular dynamics (MD) simulations using GROMACS (v.2020.1) (80). Set-up, energy minimisation, equilibration, and production simulations were performed as previously described (81), specific details of which have been provided in Supplementary methods 3. Production simulations were performed for 100 ns and the resulting trajectories were visualised. In addition, the RMSD of the position of backbone atoms was analysed as a function of simulation time using GROMACS modules gms rms and gmx gyrate respectively. The Visual Molecular Dynamics (VMD, v.1.9.4) program was used to visualise simulations and the Gnuplot (v.5.2) program was employed to produce the graphics associated with this work.

#### 4.5.4. Interface analysis

The PDBePISA web server (82) was used to analyse the interfacial interactions of modelled Cry48-Tpp49 complexes. PISA enabled calculation of the interface area (Å^2^) and Δ^i^G (kcal mol^1^), as well as the identification of interfacial hydrogen bonds and salt bridges.

## Acknowledgments

We acknowledge European XFEL in Schenefeld, Germany, for provision of X-ray free-electron laser beamtime at the SPB/SFX Instrument and use of the XBI laboratories and would like to thank the staff for their assistance. We acknowledge the support of the Supercomputing Wales project, which is part-funded by the European Regional Development Fund (ERDF) via the Welsh Government. This research was supported in part through the Maxwell computational resources operated at Deutsches Elektronen-Synchrotron DESY, Hamburg, Germany. We would like to thank Frank Schluenzen (DESY) for installation and testing of Alphafold2 on the Maxwell-cluster. We would like to thank Dr. Esther Schnettler for kindly providing us with the *C. tarsalis* (Ct) cell line. We thank Benjamin Nyman and Alec Gerry of the University of California, Riverside for the *Ae. albopictus* larvae, Jennifer Henke of the Coachella Valley Mosquito and Vector Control District for the *C. tarsalis* larvae, and Anthony James of the University of California, Irvine, for the *An. stephensi* larvae.

## Funding

This work was supported by the Biotechnology and Biological Sciences Research Council (BBSRC, grant reference BB/S002774/1) and a BBSRC-funded South West Biosciences Doctoral Training Partnership (training grant reference BB/M009122/1). PLX thanks Joachim Herz Stiftung for a fellowship. BF acknowledges partial funding support of the bioassays from the Pacific Southwest Regional Center of Excellence for Vector-Borne Diseases funded by the U.S. Centers for Disease Control and Prevention (Cooperative Agreement 1U01CK000516).

## Notes

### Competing Interest Statement

The authors have declared no competing interest.

### Summary of Updates

This version of the manuscript has been updated to include bioassays confirming toxicity against Culex tarsalis, Anopheles stephensi, and Aedes albopictus mosquito larvae. In addition, two new Tpp49Aa1 structures obtained from complementary crystallography experiments conducted at pH 3 and pH 11 shed light on the mechanisms of crystal dissolution.

