## Supplementary Information for "Structure of the *Lysinibacillus sphaericus* Tpp49Aa1 pesticidal protein elucidated from natural crystals using MHz-SFX"

**Affiliations**

**This PDF file includes:**

Figures S1 to S9

Tables S1 to S5

Supplementary methods 1

Supplementary methods 2

Supplementary methods 3

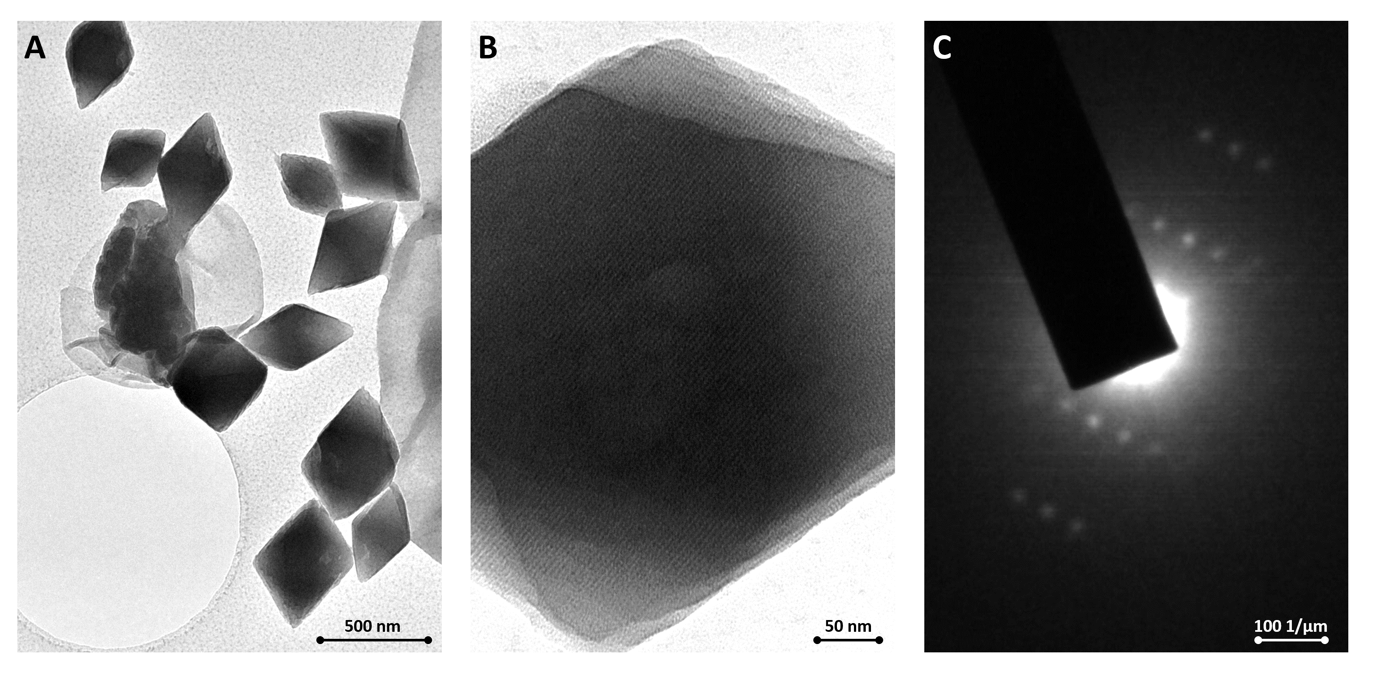

**Figure S1. Transmission electron microscopy on negative stained Tpp49Aa1 nanocrystals. (A)** Tpp49Aa1 nanocrystals and remaining parasporal bodies from crystal purification process. **(B)** Visualization of crystal lattice of Tpp49Aa1 nanocrystals allows assessment of the quality of different purification batches. **(C)** Confirmation of crystal quality is obtained by selected area electron diffraction imaging (SAED).

**Table S1. Data collection and refinement statistics for Tpp49Aa1**

| **Data Collection** | | | |
| --- | --- | --- | --- |
| PDB ID | 8BEX | 8BEY | 8BEZ |
| Beamline | SPB/SFX at European XFEL | SPB/SFX at European XFEL | SPB/SFX at European XFEL |
| X-ray Energy (keV) | 9.3 | 9.3 | 9.3 |
| Wavelength (Å) | 1.33 | 1.33 | 1.33 |
| **Crystal Data (figures in brackets refer to outer resolution shell)** | | | |
| pH | 3 | 7 | 11 |
| a,b,c (Å) | 79.89, 82.84, 157.88 | 79.65, 83.11, 156.91 | 80.12, 83.22, 156.49 |
| α,β,γ (°) | 90.0, 90.0, 90.0 | 90.0, 90.0, 90.0 | 90.0, 90.0, 90.0 |
| Space group | P 2_1_ 2_1_ 2_1_ | P 2_1_ 2_1_ 2_1_ | P 2_1_ 2_1_ 2_1_ |
| Resolution (Å) | 1.78 – 36.01 | 1.62 - 24.98 | 1.75 – 36.12 |
| Outer shell | 1.78 – 1.80 | 1.62-1.64 | 1.75-1.77 |
| *R-*split (%) | 8.59 (231.27) | 6.07 (299.67) | 6.40 (177.89) |
| CC* | 0.999 (0.492) | 0.999 (0.484) | 0.999 (0.632) |
| CC1/2 | 0.997 (0.138) | 0.997 (0.133) | 0.998 (0.249) |
| I / σ(I) | 10.50 (0.81) | 10.61 (0.36) | 14.81 (0.75) |
| Completeness (%) | 99.51 (92.76) | 100 (100) | 99.82 (97.34) |
| Multiplicity | 183.70 (6.3) | 1028.30 (186.7) | 323.70 (10.6) |
| Total Measurements | 18469814 (38681) | 136507469 (1635083) | 34281129 (71761) |
| Unique Reflections | 100541 (6177) | 132750 (8756) | 105904 (6782) |
| Wilson B-factor(Å^2^) | 27.96 | 21.79 | 23.82 |
| **Refinement Statistics** | | | |
| Refined atoms | 7,234 | 7,357 | 7,324 |
| Protein atoms | 6,867 | 6,877 | 6,854 |
| Non-protein atoms | 0 | 0 | 0 |
| Water molecules | 367 | 480 | 470 |
| R-work reflections | 95,333 | 130,631 | 100,468 |
| R-free reflections | 5,033 | 6,921 | 5,313 |
| R-work/R-free (%) | 18.9 / 21.3 | 17.8 / 19.7 | 17.7 / 19.9 |
| **rms deviations (target in brackets)** | | | |
| Bond lengths (Å) | 0.011 (0.013) | 0.013 (0.013) | 0.012 (0.013) |
| Bond Angles (°) | 1.472 (1.648) | 1.446 (1.648) | 1.443 (1.648) |
| ^1^Coordinate error (Å) | 0.105 | 0.077 | 0.088 |
| Mean B value (Å^2^) | 32.724 | 31.368 | 29.318 |
| **Ramachandran Statistics (PDB Validation)** | | | |
| Favoured/allowed/Outliers | 812 / 15 / 1 | 814/ 14 / 0 | 811 / 17 / 0 |
| % | 98.1 / 1.8 / 0.1 | 98.3 / 1.7 / 0 | 98 / 2 / 0 |

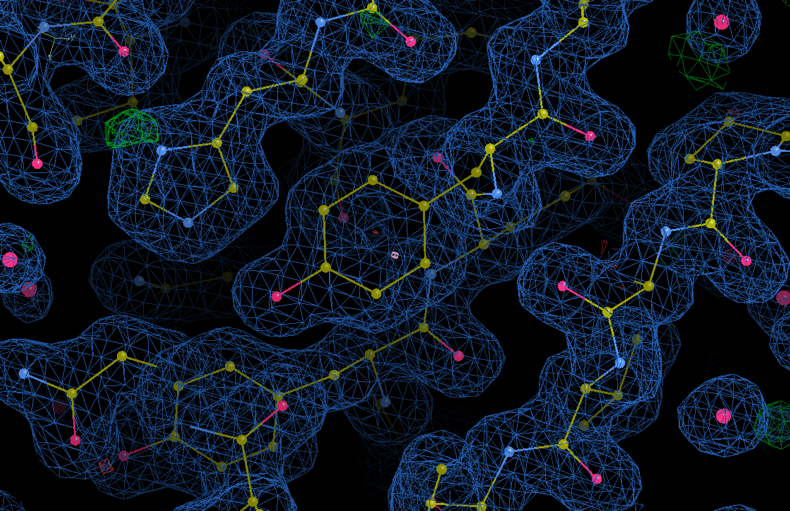

A

Tyr388

His282

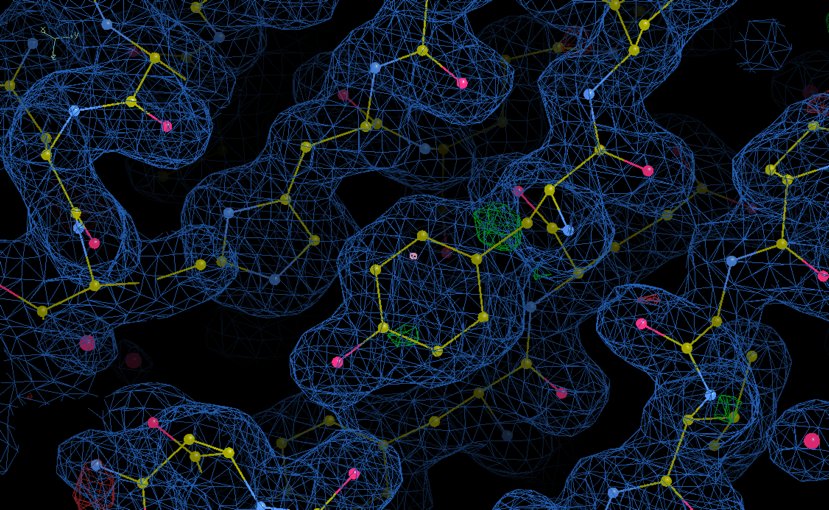

His282

Tyr388

B

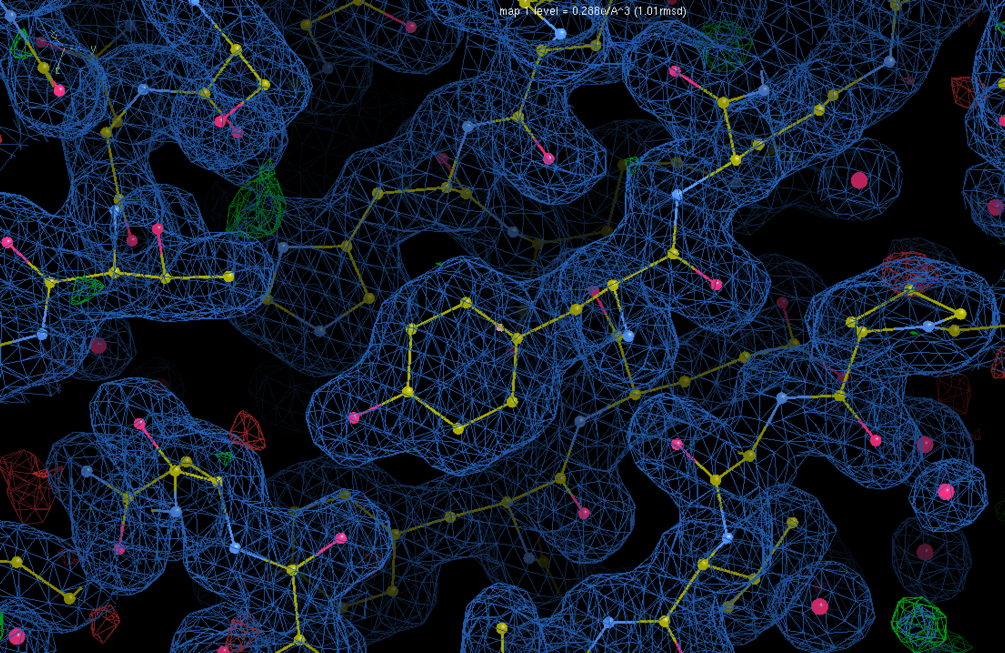

C

His282

Tyr388

**Figure S2. Electron density map and model of the XFEL structure of Tpp49Aa1.** A slice through the centre of the electron density of a monomer of Tpp49Aa1 is shown. Protein backbone is coloured yellow, nitrogen atoms are coloured blue, and oxygen atoms are coloured pink. **(A)** Tpp49Aa1, pH 7 **(B)** Tpp49Aa1, pH 3 **(C)** Tpp49Aa1, pH 11

**Table S2 – Structural similarity of Tpp49Aa1 with other β-sheet toxins. Z-scores larger than 2 are considered significant.**

| **PDB ID** | **Name (formally)** | **Z-score** | **RMSD (Å)** |
| --- | --- | --- | --- |
| 5FOY_B | Tpp2Aa2 (BinB) | 41.4 | 2.1 |
| 3WA1_A | Tpp2Aa3 (BinB) | 38.3 | 2.3 |
| 5FOY_A | Tpp1Aa2 (BinA) | 37.9 | 2.5 |
| 4JP0_A | Tpp35Ab2 (Cry35) | 29.3 | 3.5 |

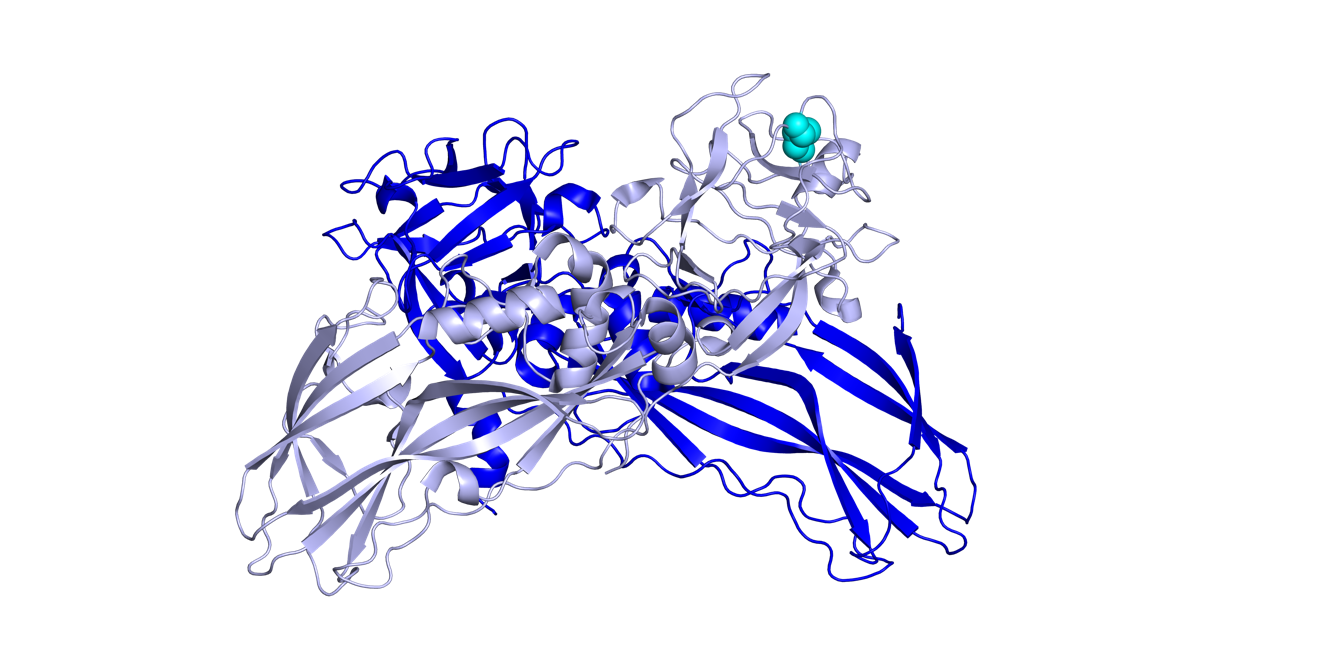

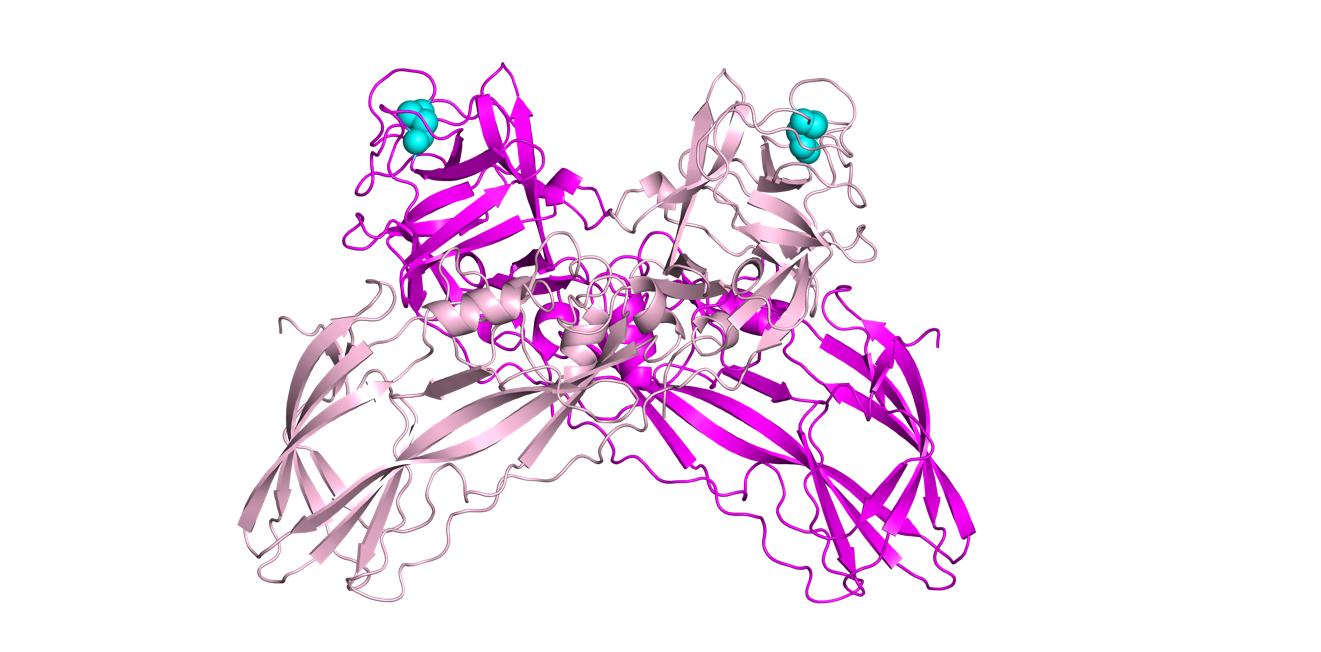

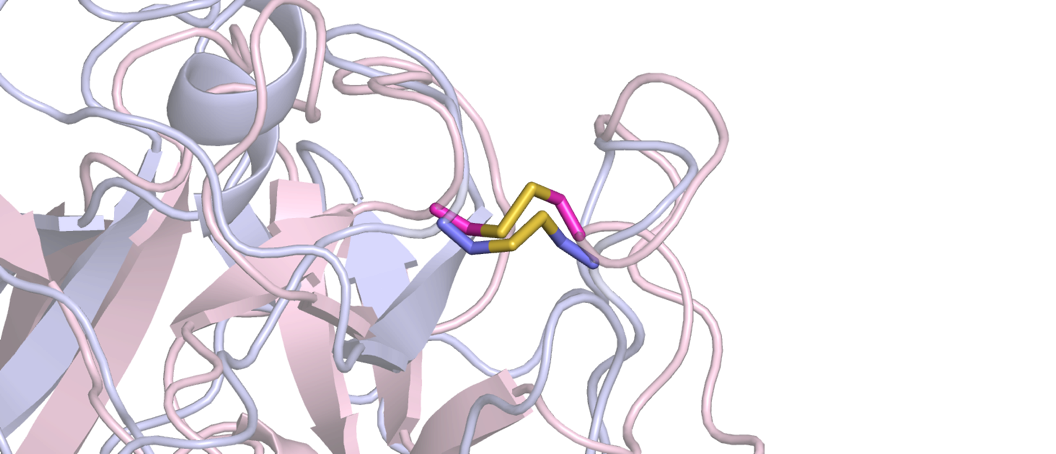

**A**

**B**

**C**

**Figure S3. Tpp49Aa1 forms a homodimer similar to the Tpp1Aa2/Tpp2Aa2 heterodimer. (A)** Tpp49Aa1 homodimer (magenta and light pink). **(B)** Tpp1Aa2 (blue)/Tpp2Aa2 (light blue) heterodimer. Equivalent disulphide bonds in Tpp49Aa1 and Tpp2Aa2 monomers are shown as spheres (cyan). In the Tpp49Aa1 homodimer, the interface between the two monomers involves 41 residues from monomer A and 42 residues from monomer B, making 16 hydrogen bonds. Interface analysis by PISA estimates the Tpp49Aa1 interface area at 1329.1 Å^2^ and the binding energy at -11.1 kcal mol^-1^. In the Tpp1Aa2/Tpp2Aa2 heterodimer, the interface between the Tpp1Aa2 and Tpp2Aa2 monomers involves 49 residues from Tpp1Aa2 and 63 residues from Tpp2Aa2, making 19 hydrogen bonds and 2 salt bridges. Consistent with this, interface analysis by PISA estimates the interface area at 1833.1 Å^2^ and the binding energy at -22.5 kcal mol^-1^, indicating a more stable complex for Tpp1Aa2/Tpp2Aa2 heterodimers than for Tpp49Aa1 homodimers. The RMSD of the aligned atoms between the Tpp49Aa1 homodimer and Tpp1Aa2/Tpp2Aa2 heterodimer was estimated by PyMOL at 9.023 Å. **(C)** Alignment of Tpp49Aa1 (light pink) and Tpp2Aa2 (light blue). Equivalent Tpp49Aa1 disulphide bond, Cys91-Cys183, and Tpp2Aa2 disulphide bond, Cys67-Cys161, are shown as sticks.

**Figure S4. Superposition of the Tpp49Aa1 monomers.** Showing the two copies to be almost identical, with an all-atom RMSD of 0.681 Å. **(A)** Superposition of Tpp49Aa1 monomers, represented as cartoon. **(B)** Superposition of Tpp49Aa1 monomers, represented by B-factor putty. The B-factor, or thermal parameter, describes the magnitude of displacement of the atoms from their central positions.

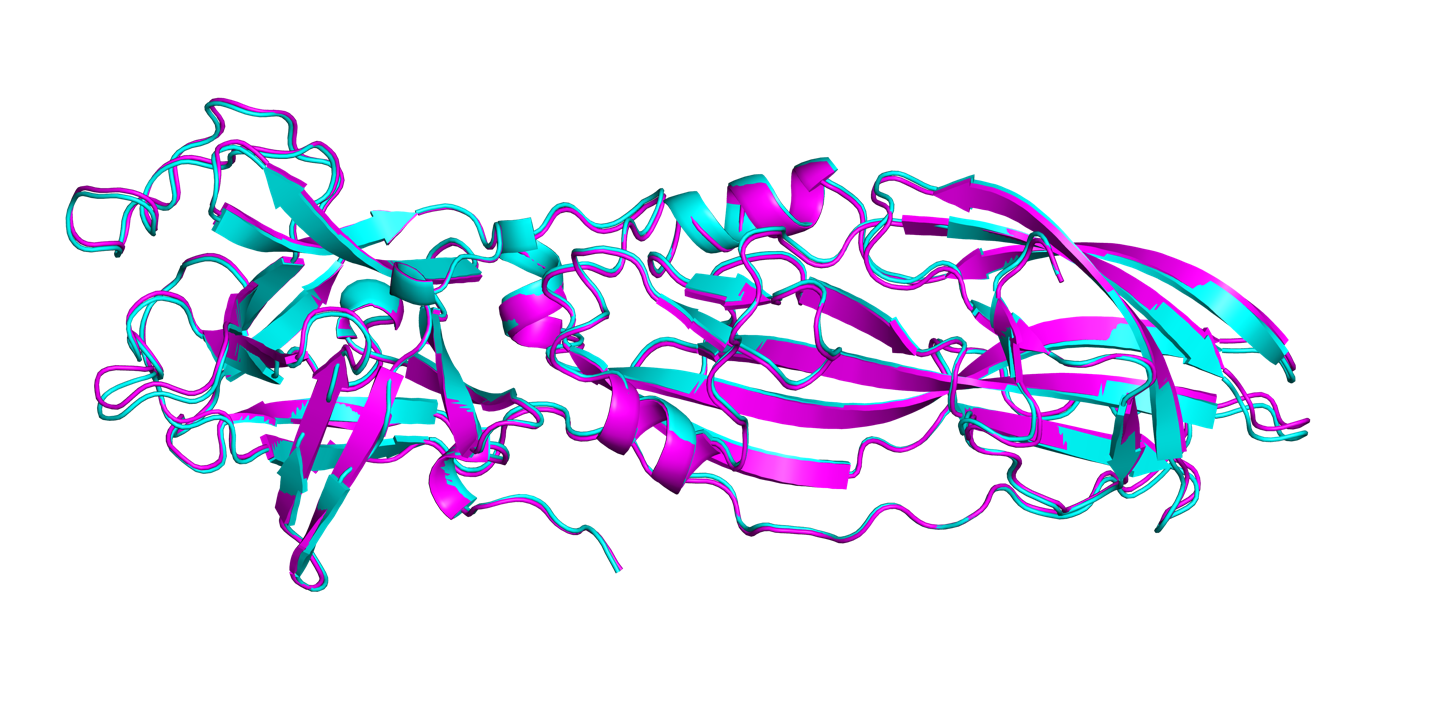

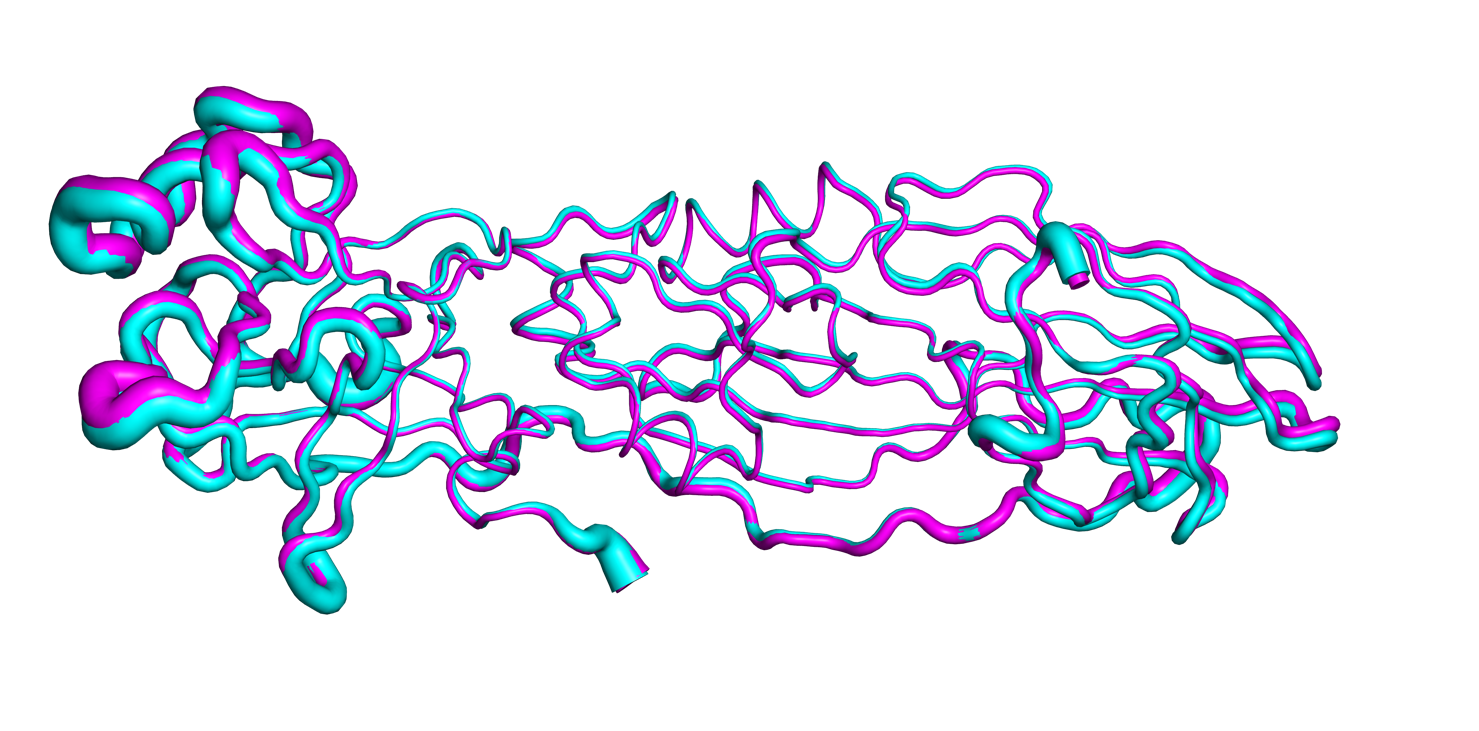

**A**

**B**

| **Interface** | **Monomer 1** | | | **Monomer 2** | | | **Interface**  **area (Å^2^)** | **Δ^i^G****  **(kcal mol^-1^)** | **H-bonds** | **Salt bridges** | **Binding energy**  **(kcal mol^-1^)** |
| --- | --- | --- | --- | --- | --- | --- | --- | --- | --- | --- | --- |
|  | **Chain** | **Symmetry** | **Nres** | **Chain** | **Symmetry** | **Nres** |  |  |  |  |  |
| 1* | B | x,y,z | 42 | A | x,y,z | 41 | 1329.1 | -4.0 | 16 | 0 | -11.1 |
| 2 | A | x,y,z | 13 | B | x-1/2,-y-1/2,-z | 13 | 470.2 | -0.6 | 10 | 2 | -5.8 |
| 3 | B | x,y,z | 12 | A | -x,y-1/2,-z-1/2 | 13 | 445.1 | -0.3 | 7 | 1 | -3.8 |
| 4 | B | x,y,z | 12 | A | -x+1/2,-y,z-1/2 | 10 | 383.4 | -8.0 | 0 | 0 | -8.0 |
| 5 | B | x,y,z | 20 | B | x-1/2,-y-1/2,-z | 15 | 361.3 | -0.1 | 5 | 0 | -2.3 |
| 6 | A | x,y,z | 12 | A | -x,y-1/2,-z-1/2 | 8 | 147.9 | 0.9 | 1 | 0 | 0.5 |
| 7 | A | x,y,z | 6 | B | x-1,y,z | 6 | 169.1 | -1.3 | 0 | 1 | -1.7 |
| 8 | A | x,y,z | 3 | B | -x-1/2,-y-1,z-1/2 | 3 | 91.6 | 0.5 | 1 | 3 | -1.0 |
| 9 | B | x,y,z | 1 | A | x,y-1,z | 2 | 12.4 | 0.3 | 0 | 0 | 0.3 |
| Total*** | – | – | – | – | – | – | 2081 | -8.6 | 24 | 7 | -21.8 |

**Table S3. Structural properties of the molecular interfaces in the Tpp49Aa1 crystal at pH 7**

**Table S4. Structural properties of the molecular interfaces in the Tpp49Aa1 crystal at pH 3**

| **Interface** | **Monomer 1** | | | **Monomer 2** | | | **Interface**  **area (Å^2^)** | **Δ^i^G****  **(kcal mol^-1^)** | **H-bonds** | **Salt bridges** | **Binding energy**  **(kcal mol^-1^)** |
| --- | --- | --- | --- | --- | --- | --- | --- | --- | --- | --- | --- |
|  | **Chain** | **Symmetry** | **Nres** | **Chain** | **Symmetry** | **Nres** |  |  |  |  |  |
| 1* | B | x,y,z | 40 | A | x,y,z | 40 | 1333.5 | -5.0 | 16 | 0 | -12.1 |
| 2 | A | x,y,z | 14 | B | x-1/2,-y-1/2,-z | 14 | 466.5 | -1.1 | 6 | 0 | -3.8 |
| 3 | B | x,y,z | 12 | A | -x,y-1/2,-z-1/2 | 13 | 445.0 | -0.4 | 9 | 0 | -4.4 |
| 4 | B | x,y,z | 10 | A | -x+1/2,-y,z-1/2 | 11 | 354.0 | -7.7 | 0 | 0 | -7.7 |
| 5 | B | x,y,z | 18 | B | x-1/2,-y-1/2,-z | 11 | 305.2 | 1.9 | 3 | 0 | 0.5 |
| 6 | A | x,y,z | 11 | A | -x,y-1/2,-z-1/2 | 6 | 145.4 | 1.1 | 1 | 0 | 0.6 |
| 7 | A | x,y,z | 6 | B | x-1,y,z | 5 | 113.7 | -1.0 | 0 | 1 | -1.3 |
| 8 | A | x,y,z | 3 | B | -x-1/2,-y-1,z-1/2 | 3 | 89.2 | 1.1 | 1 | 4 | -0.9 |
| 9 | B | x,y,z | 1 | A | x,y-1,z | 2 | 7.1 | 0.2 | 0 | 0 | 0.2 |
| Total*** | – | – | – | – | – | – | 1926.1 | -5.9 | 20 | 5 | -16.8 |

**Table S5. Structural properties of the molecular interfaces in the Tpp49Aa1 crystal at pH 11**

| **Interface** | **Monomer 1** | | | **Monomer 2** | | | **Interface**  **area (Å^2^)** | **Δ^i^G****  **(kcal mol^-1^)** | **H-bonds** | **Salt bridges** | **Binding energy**  **(kcal mol^-1^)** |
| --- | --- | --- | --- | --- | --- | --- | --- | --- | --- | --- | --- |
|  | **Chain** | **Symmetry** | **Nres** | **Chain** | **Symmetry** | **Nres** |  |  |  |  |  |
| 1* | B | x,y,z | 38 | A | x,y,z | 41 | 1312.8 | -5.2 | 15 | 0 | -11.9 |
| 2 | A | x,y,z | 13 | B | x-1/2,-y-1/2,-z | 13 | 477.8 | 0.2 | 9 | 1 | -4.2 |
| 3 | B | x,y,z | 13 | A | -x,y-1/2,-z-1/2 | 13 | 458.3 | -1.0 | 7 | 1 | -4.5 |
| 4 | B | x,y,z | 12 | A | -x+1/2,-y,z-1/2 | 11 | 396.4 | -7.0 | 0 | 0 | -7.0 |
| 5 | B | x,y,z | 19 | B | x-1/2,-y-1/2,-z | 15 | 355.3 | -0.5 | 4 | 0 | -2.3 |
| 6 | A | x,y,z | 10 | A | -x,y-1/2,-z-1/2 | 7 | 131.5 | 0.9 | 1 | 0 | 0.5 |
| 7 | A | x,y,z | 6 | B | x-1,y,z | 6 | 180.1 | -1.4 | 0 | 1 | -1.7 |
| 8 | A | x,y,z | 3 | B | -x-1/2,-y-1,z-1/2 | 3 | 89.6 | 0.5 | 1 | 3 | -1.1 |
| 9 | B | x,y,z | 2 | A | x,y-1,z | 3 | 12.8 | 0.3 | 0 | 0 | 0.3 |
| Total*** | – | – | – | – | – | – | 2101.8 | -8 | 22 | 6 | -20 |

* Interface 1 refers to the Tpp49Aa1 dimer interface

** Δ^i^G = solvation free energy gain

*** Total excludes dimer interface

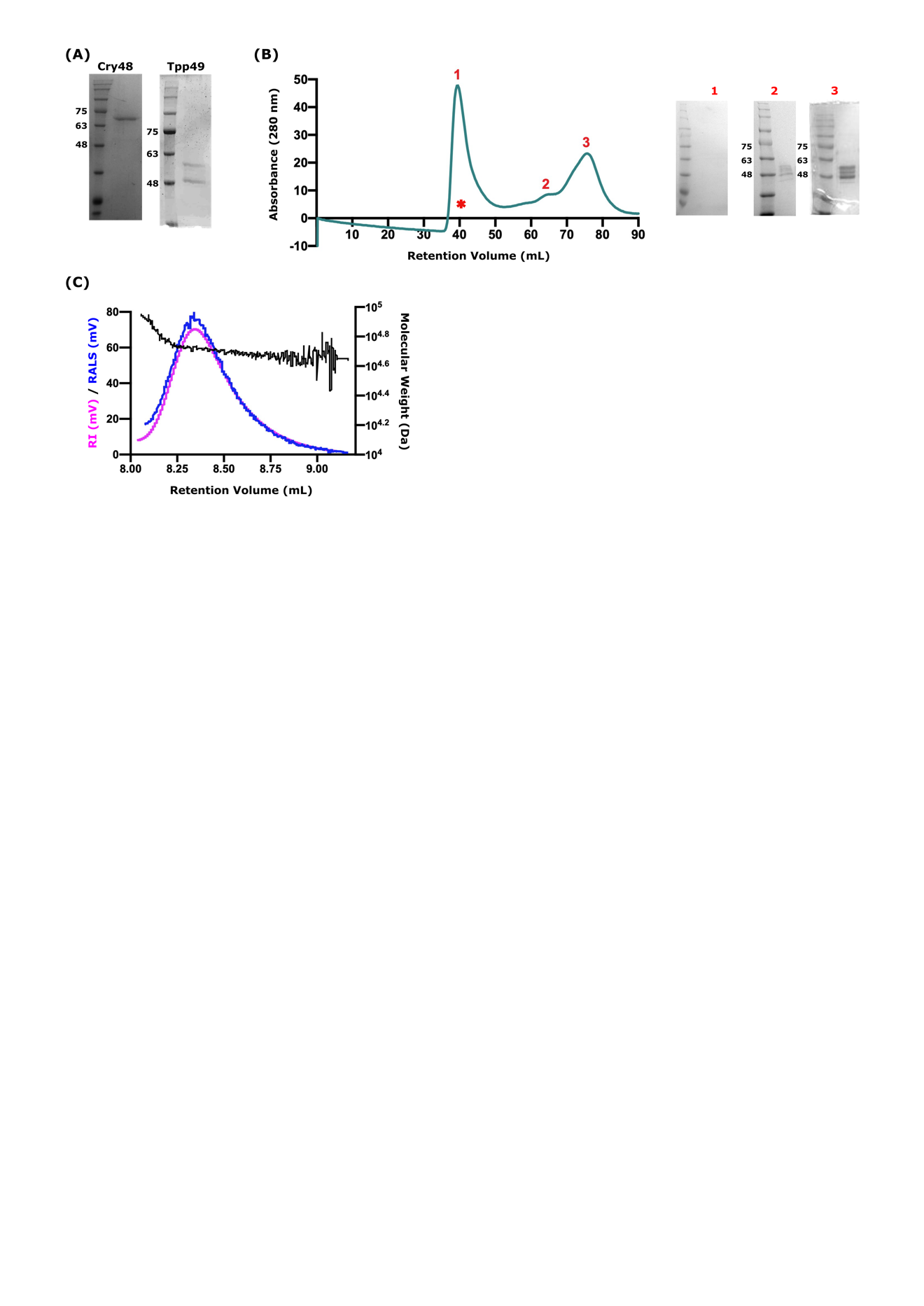

**Figure S5. Tpp49Aa1 is predominantly monomeric in solution.** A combination of size exclusion chromatography (SEC) and static light scattering (RALS) was used to determine if Tpp49Aa1 is dimeric or monomeric in solution. (**A)** Crystalline protein solubilised in Na_2_CO_3_ overnight showing Cry48Aa1 at ~ 70 kDa and Tpp49Aa1 as two bands at ~ 49 and ~55 kDa. (**B**) SEC shows three absorbance peaks (UV 280 nm) representing 766, 121, and 51 kDa, respectively. SDS-page resolution of the fractions taken from those peaks shows no product at 1 (*column void volume, presumably non-protein contaminants), and bands of approximately the expected sizes in 2 and 3. **(C)** Protein-containing peaks (2 & 3) were concentrated and used to decipher molecular weight via Right Angle Light Scattering (RALS, blue) and Refractive Index (RI, pink) measurements. From these data the molecular weight (Da, black) for the protein was calculated to be 52.1 kDa when calibrated to BSA (1mg/ mL) using OmniSEC software.

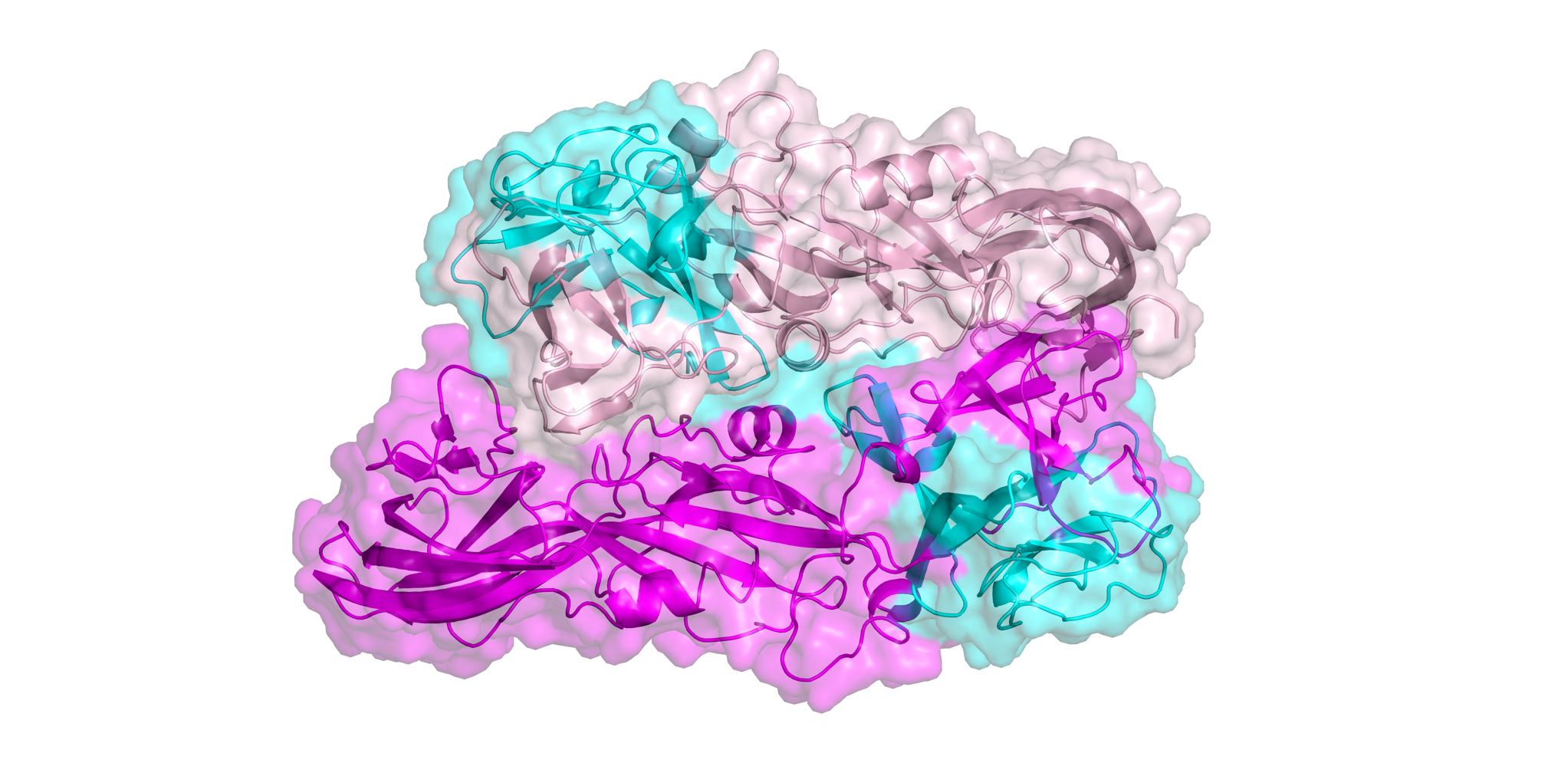

**Figure S6. Tpp49Aa1 dimer highlighting the regions proposed to interact with Cry48Aa1.** Tpp49Aa1 regions proposed to interact with Cry48Aa1 (cyan) are partially buried within the dimer interface. Tpp49Aa1 monomers shown in magenta and pink.

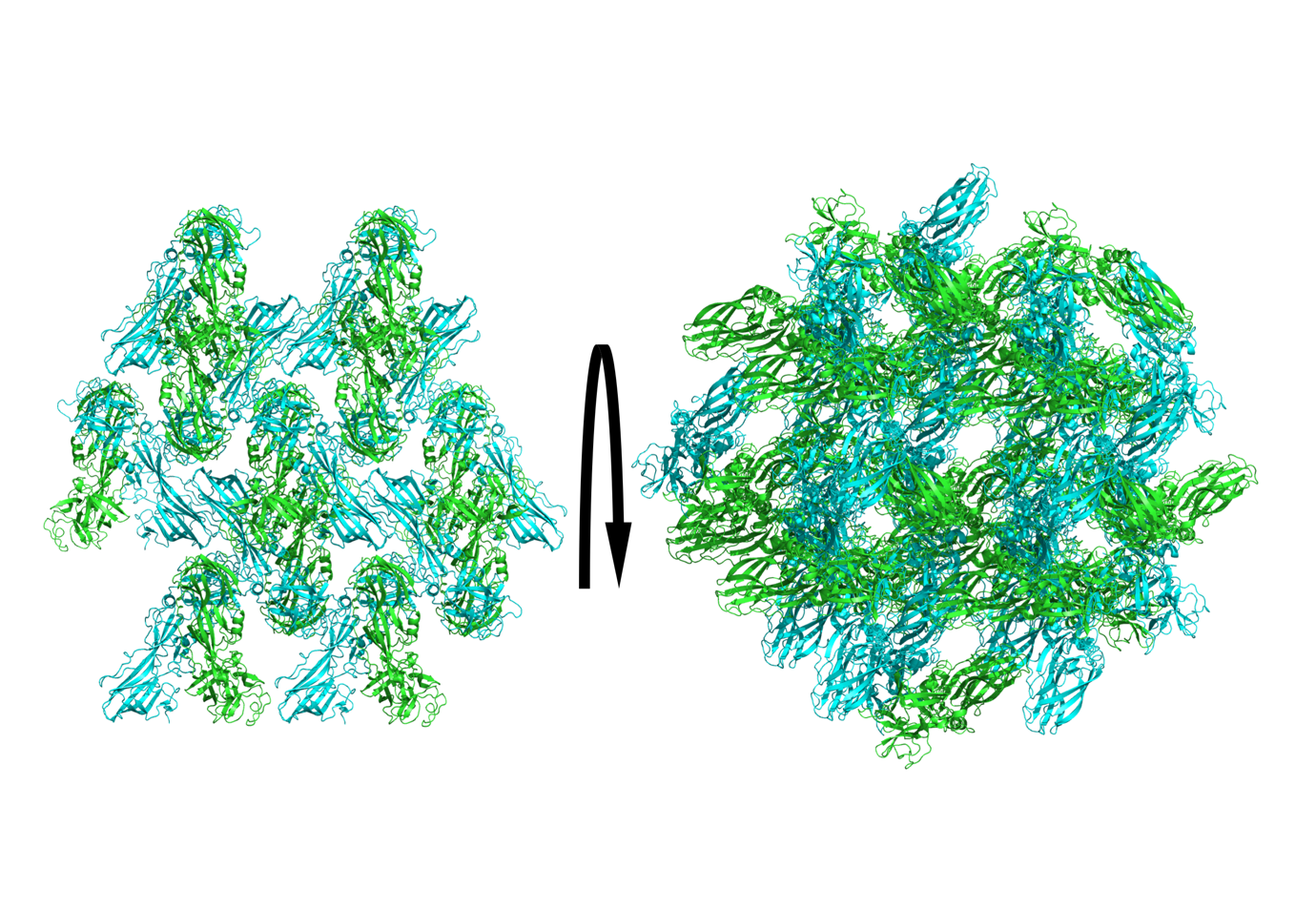

**Figure S7. Solvent channels in Tpp49Aa1 crystals**

**Table S6A. Loss/formation of interactions perturbed by an increase in pH from 7 to 11**

| **Interface** | Monomer 1 - symmetry | Monomer 2 - symmetry | Interactions |
| --- | --- | --- | --- |
| 1* | x,y,z | x,y,z | Loss of:   - B:Asn51(ND2) – A:Asp347(O) - B:Asn51(ND2) – A:Asn350(OD1) - B:Gln162(OE1) – A:Asn357(ND2) - B:Asn347(O) – A:Asn51(ND2) - B:Asn350(OD1) – A:Asn51(ND2) - B:Thr351(N) – A:Asn51(OD1)   Formation of:   - B:Asn51(ND2) – A:Thr351(O) - B:Asp120(O) – A:Gln343(NE2) - B:Gln343(NE2) – A:Asp120(O) - B:Thr351(O) – A:Asn51(ND2) - B:Tyr435(OH) – A:Asn165(O) |
| 2 | x,y,z | x-1/2,-y-1/2,-z | Loss of:   - A:Asp429(OD1) – B:Arg267(NH1) ** - A:Tyr463(O) – B:Thr400(OG1) |
| 3 | x,y,z | -x,y-1/2,-z-1/2 | Loss of:   - B:Tyr463(O) – A:Thr400(OG1)   Formation of:   - B:Asp432(O) – A:Gln274(NE2) |
| 4 | x,y,z | -x+1/2,-y,z-1/2 |  |
| 5 | x,y,z | x-1/2,-y-1/2,-z | Loss of:   - B:Asp198(OD1) – B:Gln220(NE2) |
| 6 | x,y,z | -x,y-1/2,-z-1/2 |  |
| 7 | x,y,z | x-1,y,z | Loss of:   - A:His97(ND1) – B:Glu237(OE1) |
| 8 | x,y,z | -x-1/2,-y-1,z-1/2 | Loss of:   - A:Asp88(N) – B:Asp88(OD1) |
| 9 | x,y,z | x,y-1,z |  |

* Interface 1 refers to the Tpp49Aa1 dimer interface

** Salt bridges

**Table S6B. Loss/formation of interactions perturbed by a decrease in pH from 7 to 3**

| **Interface** | **Monomer 1 - symmetry** | **Monomer 2 - symmetry** | **Interactions** |
| --- | --- | --- | --- |
| 1* | x,y,z | x,y,z | Loss of:   - B:Gln162(OE1) – A:Asn357(ND2)   Formation of:   - B:Asn51(OD1) – A:Thr351(N) |
| 2 | x,y,z | x-1/2,-y-1/2,-z | Loss of:   - A:Asp429(OD1) – B:Arg267(NH1) ** - A:Asp429(OD2) – B:Arg267(NH1) ** - A:Ser434(N) – B:Gln274(OE1) - A:Asp429(OD2) – B:Arg267(NH1) - A:Asp429(O) – B:Arg267(NH1) - A:Asp432(O) – B:Gln274(NE2) - A:Tyr463(O) – B:Thr400(OG1) |
| 3 | x,y,z | -x,y-1/2,-z-1/2 | Loss of:   - B:Asp429(OD2) – A:Arg267(NH2) ** - B:Asp429(OD2) – A:Arg267(NH2)   Formation of:   - B:Ser434(N) – A:Asp394(OD2) - B:Asp432(O) – A:Gln274(NE2) |
| 4 | x,y,z | -x+1/2,-y,z-1/2 |  |
| 5 | x,y,z | x-1/2,-y-1/2,-z | Loss of:   - B:Arg196(NH2) – B:Gln220(OE1) - B:Asn204(ND2) – B:Gln220(O) - B:Asn206(ND2) – B:Asp394(O)   Formation of:   - B:Asn204(ND2) – B:Gln220(OE1) |
| 6 | x,y,z | -x,y-1/2,-z-1/2 |  |
| 7 | x,y,z | x-1,y,z | Loss of:   - A:His97(ND1) – B:Glu237(OE1) ** - A:His96(ND1) – B:Glu237(OE1)   Formation of:   - A:His97(NE2) – B:Glu237(OE2) ** |
| 8 | x,y,z | -x-1/2,-y-1,z-1/2 | Loss of:   - A:Asp88(N) – B:Asp88(OD1)   Formation of:   - A:Asp88(OD2) – B:Arg92(NE) ** - A:Asp88(N) – B:Asp88(OD2) |
| 9 | x,y,z | x,y-1,z |  |

* Interface 1 refers to the Tpp49Aa1 dimer interface

** Salt bridges

**Figure S8. Tpp49Aa1 and Cry48Aa1 Alphafold prediction. (A)** The suitability of the Alphafold2 package to predict the structures of pesticidal proteins was assessed by predicting the Tpp49Aa1 structure elucidated here. The Alphafold Tpp49Aa1 model (grey) displayed an all-atom RMSD of 1.982 Å (calculated using Pymol) when superposed with the Tpp49Aa1 structure (cyan). **(B)** The Tpp49Aa1 and **(C)** Cry48Aa1 Alphafold2 prediction coloured by the per-residue confidence estimate (predicted local-distance difference test - pLDDT). Model confidence scores are colour coded as follows: red – very high (pLDDT > 90), orange – confident (90 > pLDDT > 70), light blue – low (70 > pLDDT > 50), dark blue – very low (pLDDT < 50).

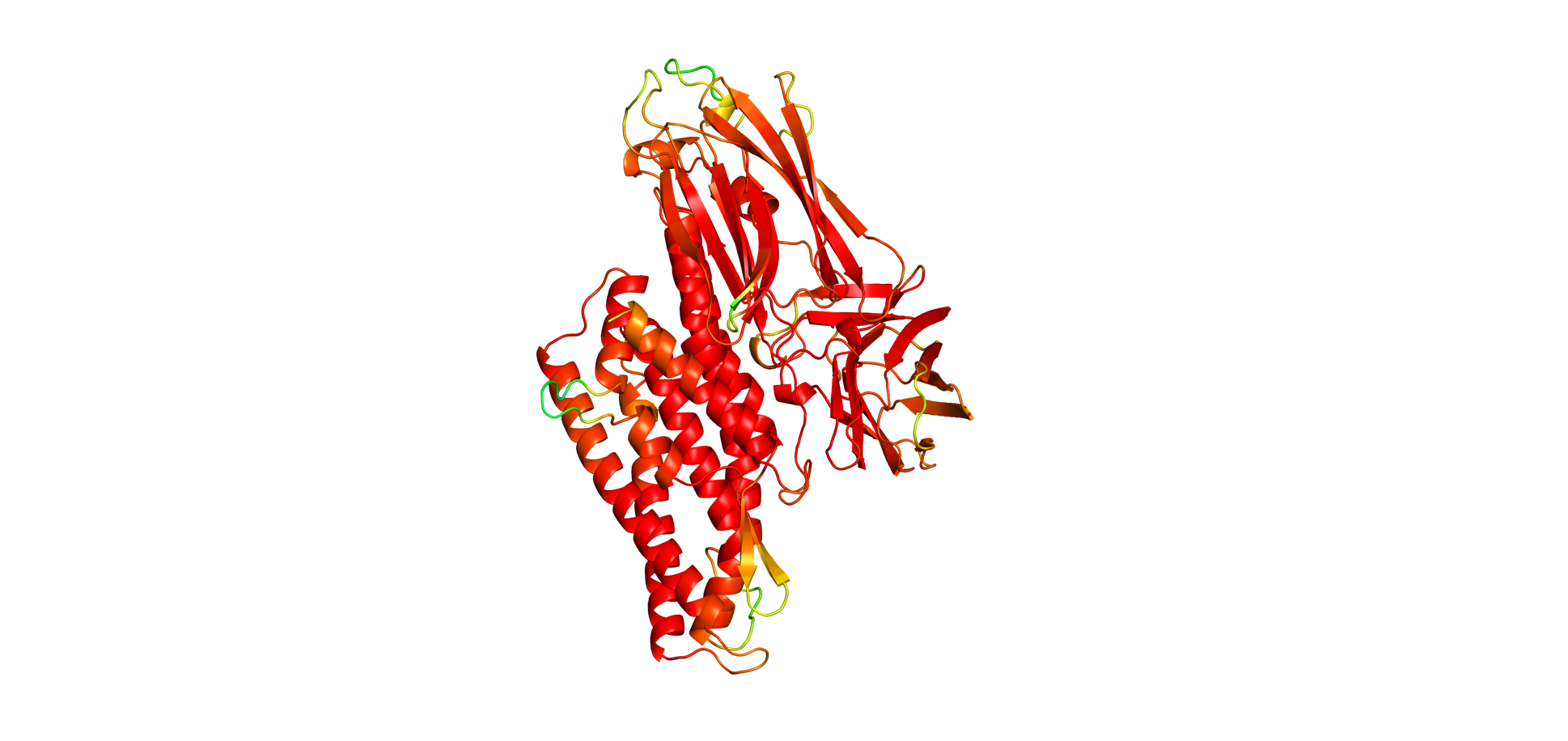

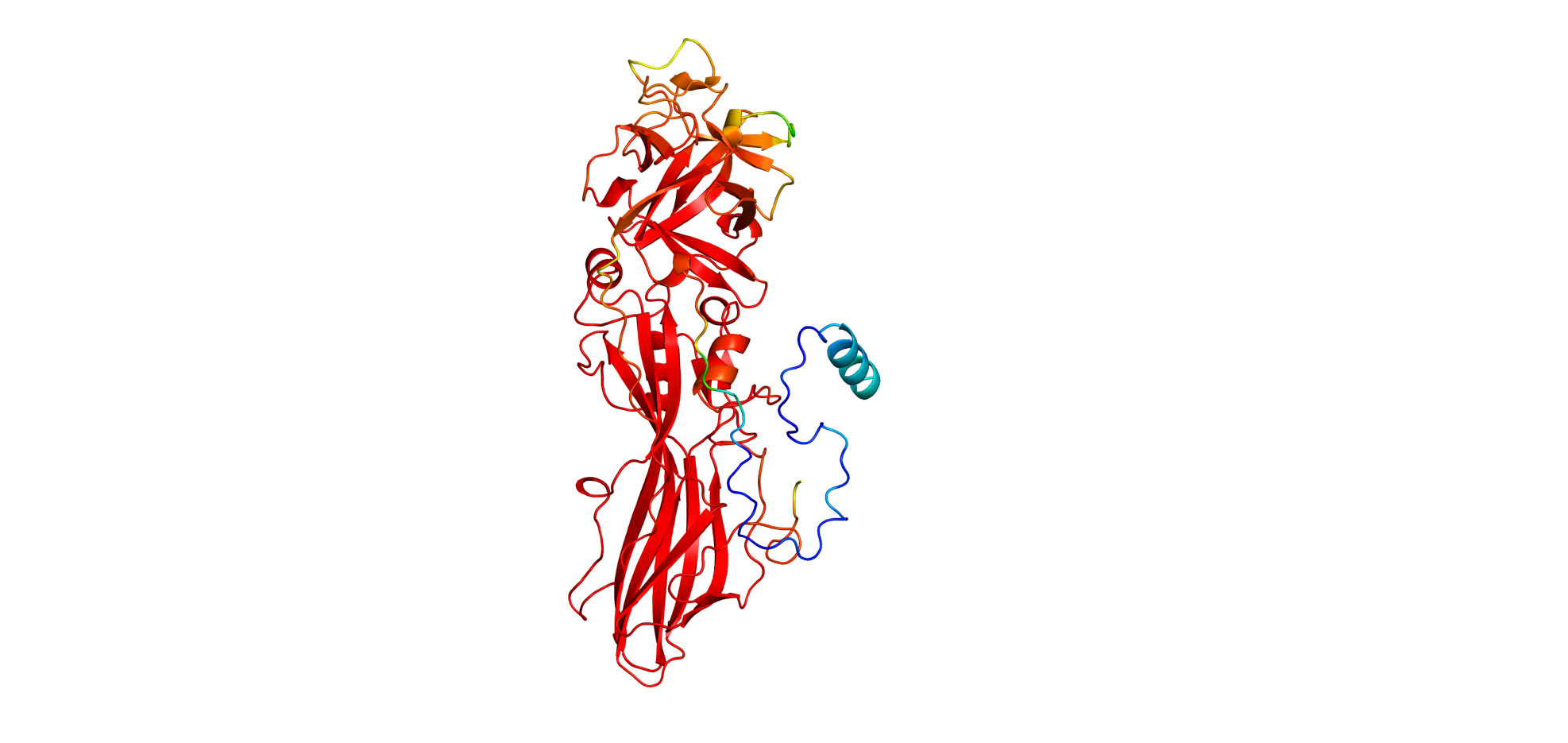

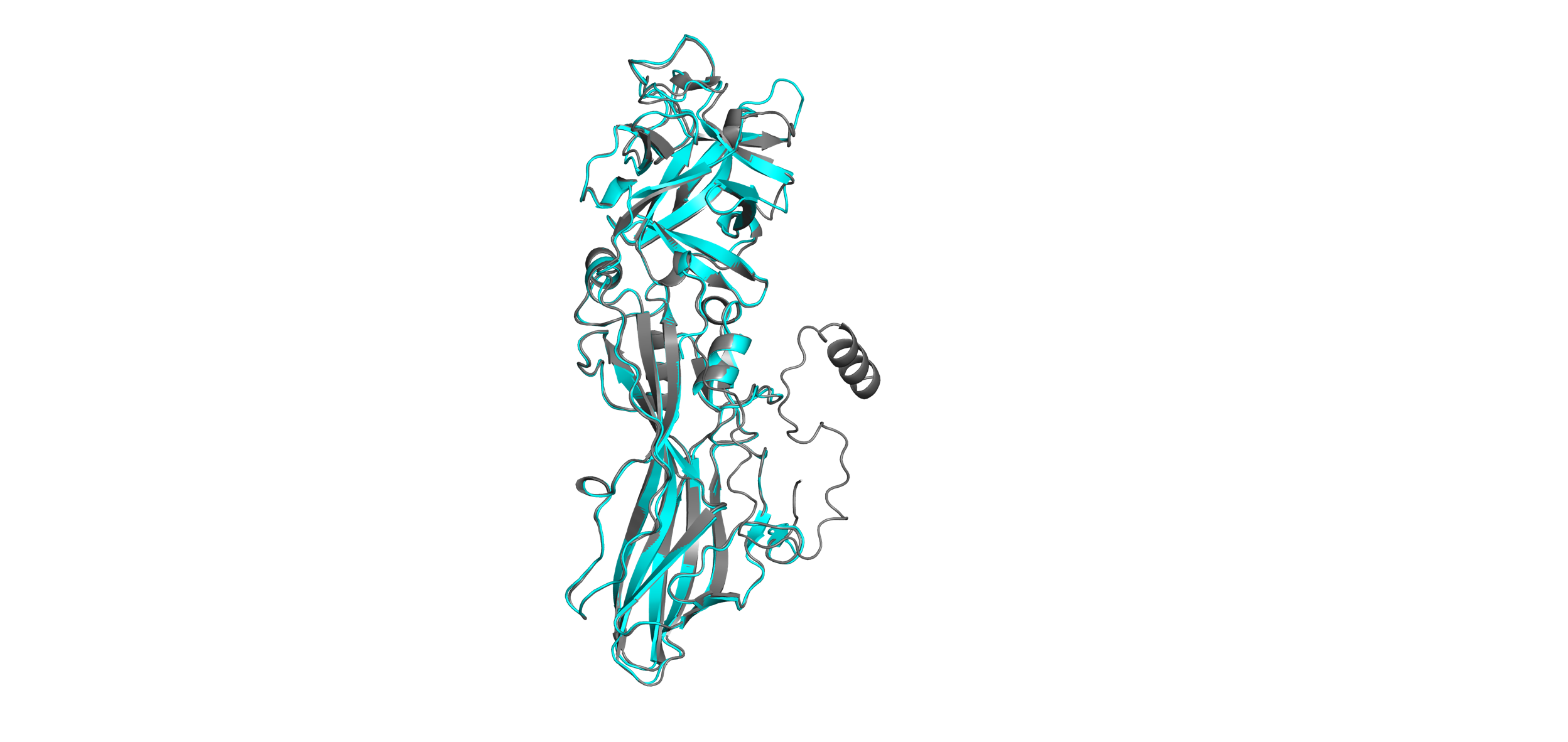

**A**

**B**

**C**

**Table S7. Computed energy and interfacial interactions of Cry48Aa1-Tpp49Aa1 modelled complexes**

| **Model** | **Energy (REU*)** | **Interface area (Å^2^)** | [**Δ^i^G**](javascript:openWindow('pi_ilist_ipv.html',400,250);) **(kcal mol^-1^)** | **H-bonds** | **Salt bridges** |
| --- | --- | --- | --- | --- | --- |
| c1r1 | -1689.582 | 907.4 | -7.9 | 2 | 0 |
| c1r2 | -1689.438 | 955.0 | -6.3 | 2 | 0 |
| c1r3 | -1689.133 | 718.7 | -5.3 | 4 | 0 |
| c1r4 | -1689.051 | 826.0 | -5.5 | 3 | 0 |
| c1r5 | -1688.059 | 810.4 | -5.0 | 2 | 0 |
| c2r1 | -1615.144 | 479.9 | -5.6 | 0 | 0 |
| c2r2 | -1612.995 | 478.8 | -5.9 | 0 | 0 |
| c2r3 | -1612.868 | 483.8 | -6.1 | 1 | 0 |
| c2r4 | -1612.415 | 531.4 | -6.4 | 2 | 0 |
| c2r5 | -1612.410 | 503.7 | -4.3 | 4 | 0 |
| c3r1 | -1696.855 | 1123.5 | -4.4 | 8 | 0 |
| c3r2 | -1696.800 | 1052.4 | -5.3 | 7 | 0 |
| c3r3 | -1695.430 | 1066.6 | -4.3 | 7 | 0 |
| c3r4 | -1695.278 | 1076.4 | -5.2 | 7 | 0 |
| c3r5 | -1695.196 | 1101.1 | -5.4 | 6 | 0 |
| c4r1 | -1583.548 | 735.6 | -6.7 | 3 | 0 |
| c4r2 | -1582.840 | 339.8 | -3.1 | 2 | 0 |
| c4r3 | -1581.636 | 802.1 | -7.4 | 2 | 0 |
| c4r4 | -1581.240 | 736.8 | -5.1 | 4 | 1 |
| c4r5 | -1580.964 | 524.9 | -5.9 | 1 | 0 |
| c5r1 | -1730.571 | 674.7 | -6.5 | 4 | 0 |
| c5r2 | -1729.703 | 534.0 | -4.4 | 3 | 0 |
| c5r3 | -1727.973 | 682.3 | -6.8 | 3 | 0 |
| c5r4 | -1727.945 | 572.9 | -4.8 | 3 | 2 |
| c5r5 | -1727.607 | 558.2 | -4.2 | 4 | 0 |

*REU = Rosetta energy units

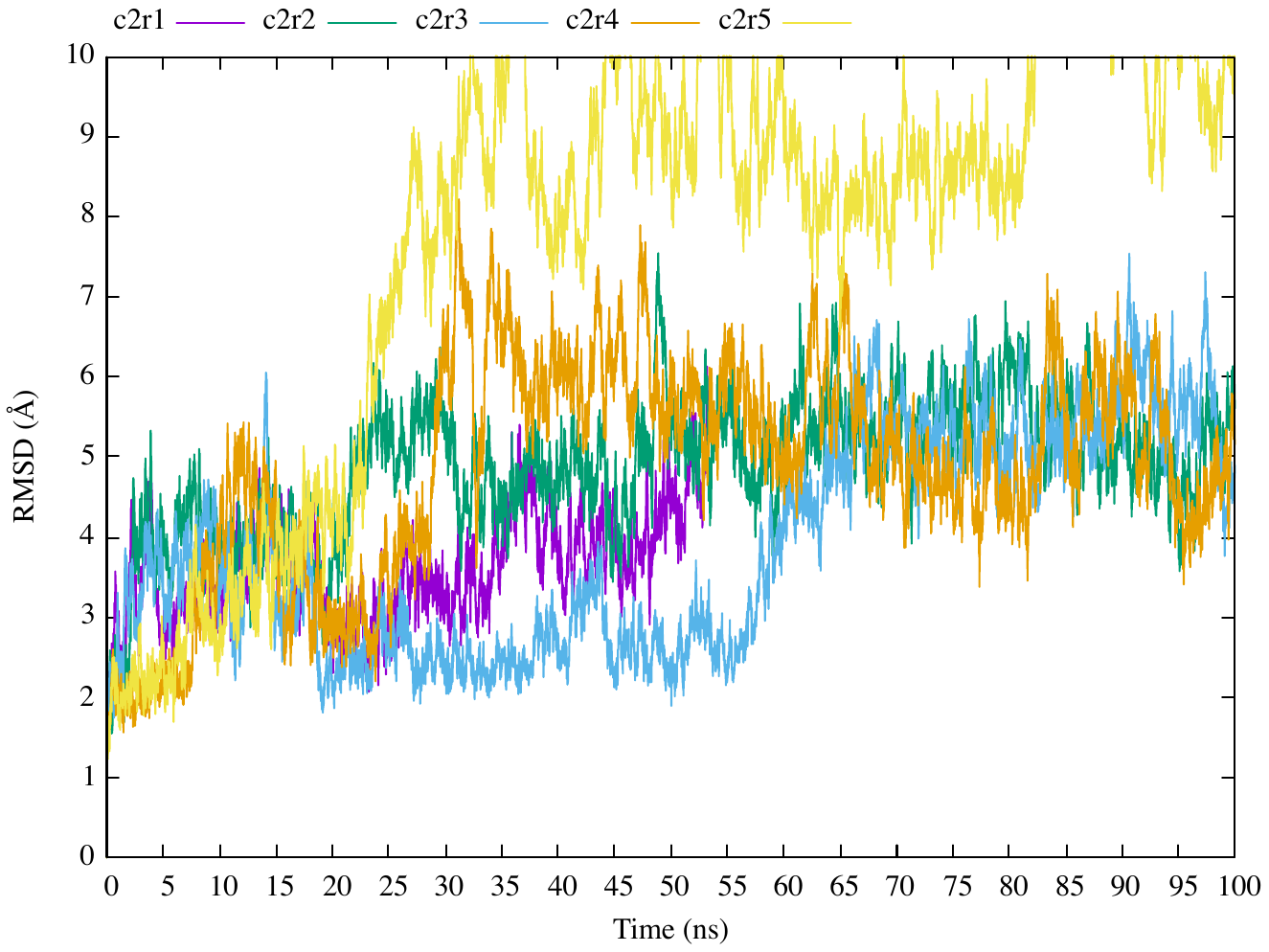

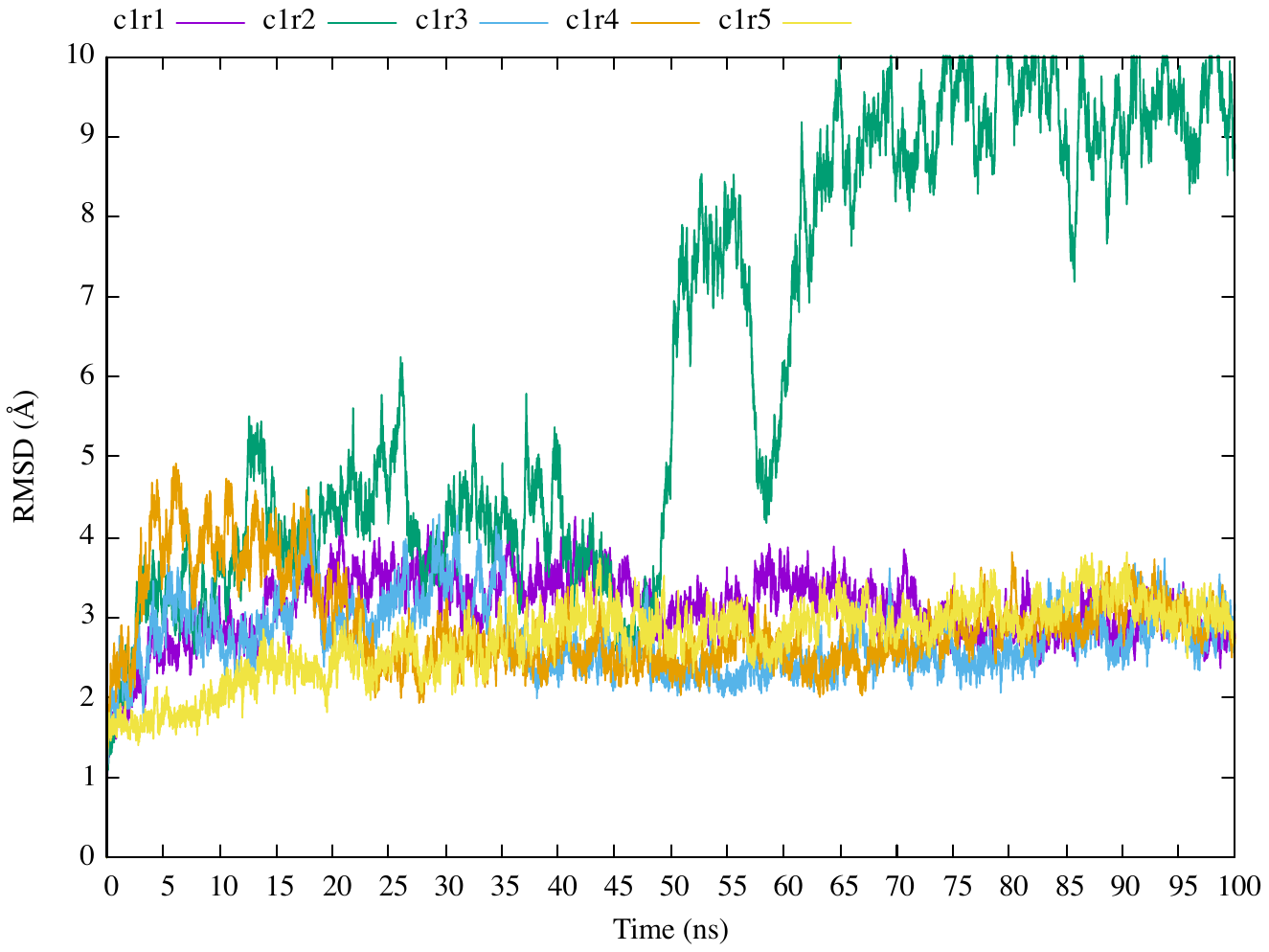

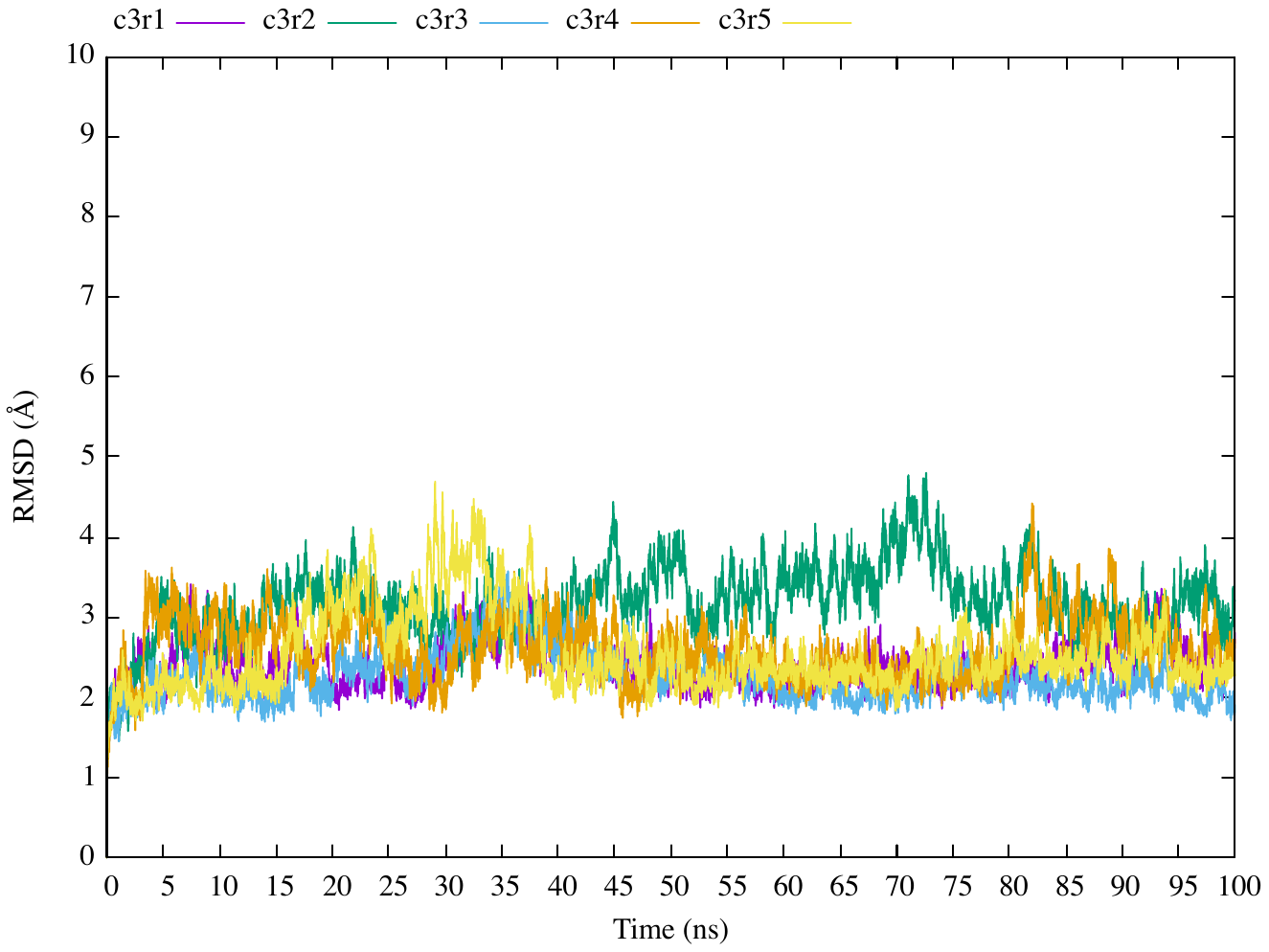

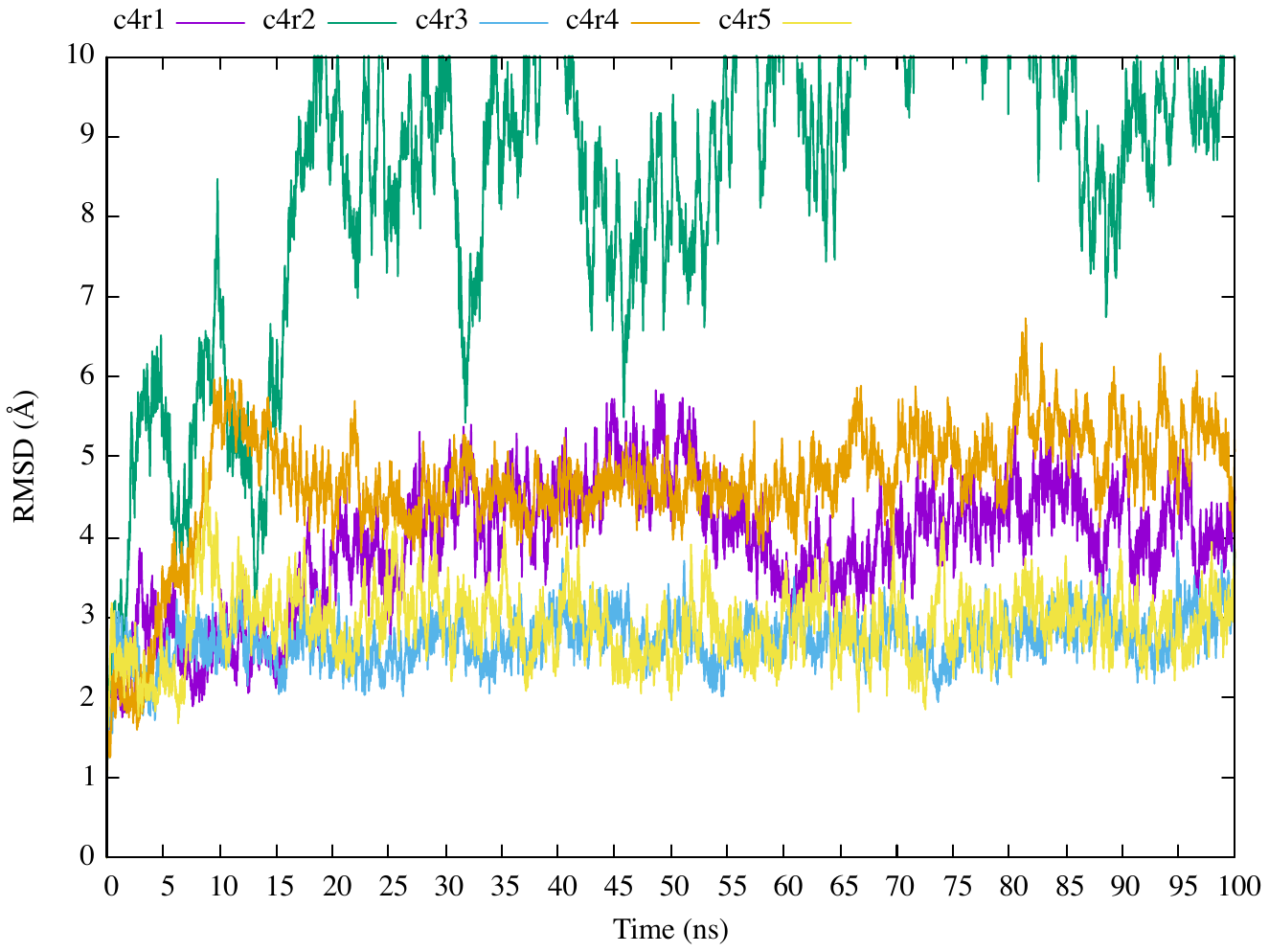

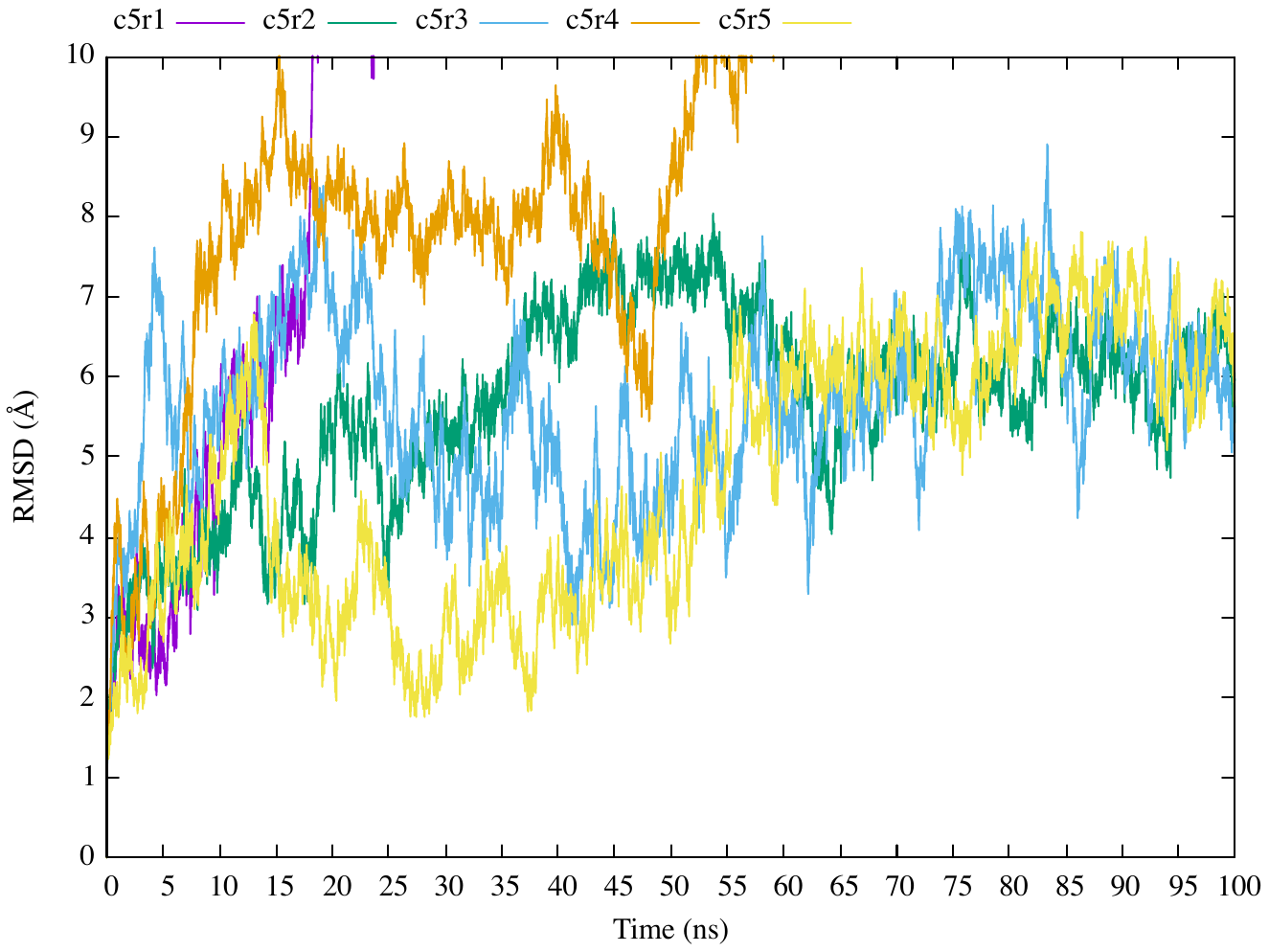

**A**

**B**

**C**

**D**

**E**

**Figure S9. Molecular dynamics simulations of Cry48Aa1-Tpp49Aa1 modelled complexes.** Root-mean-square deviation of the position of backbone atoms relative to the starting structure in Cry48Aa1-Tpp49Aa1 modelled complexes throughout 100 ns molecular dynamics simulations. Models are referred to according to their ClusPro cluster (c1 – c5) followed by their Rosetta energy rank (r1 – r5, with r1 being the lowest energy score and r5 being the highest energy score). **(A)** Cluster 1 **(B)** Cluster 2 **(C)** Cluster 3 **(D)** Cluster 4 **(E)** Cluster 5. The Gnuplot (v.5.2) program was employed to produce the graphics associated with this work.

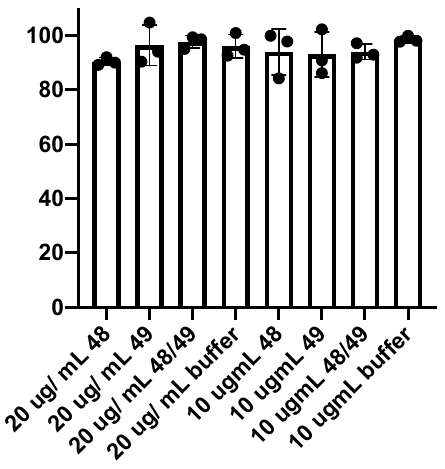

**Cell Viability**

**(% of control)**

**Figure S10. SF9 cells show no reduction in viability after 48-hour exposure to Tpp49Aa1/Cry48Aa1.** Resazurin was used to quantify the effect of these proteins on cell viability. SF9 cells were treated with a range of concentrations of either Cry48Aa1 (“48”), Tpp49Aa1 (“49”), or equimolar amounts in combination (48/49). Resazurin was added to the cells 48 h post toxin exposure. No significant difference was observed. All data are presented as percentage of control conditions (buffer only) with the mean ± SD and all statistical analysis was performed using a one-way ANOVA.

**Supplementary methods 1 - Size Exclusion Chromatography**

For size exclusion chromatography (SEC), Cry48Aa1 and Tpp49Aa1 protein crystals were solubilised in 50 mM Na_2_CO_3_ pH 10.5 + 0.05% β-mercaptoethanol overnight at room temperature, with agitation. Insoluble material was removed by centrifugation, and the solubilised proteins buffer exchanged into 20 mM TrisHCl pH 8.5. SEC was performed using a calibrated Hiload^TM^ 16/60 Superdex^TM^ S200 pg column. For column calibration, BioRad standard proteins (of molecular weights 670, 158, 44, 17, and 1.35 kDa) were run on the column at a flow rate of 0.5 mL/min with 50 mM TrisHCl buffer pH 8. All protein samples were run on the column at a flow rate of 1 mL/min with 20 mM TrisHCl pH 8.5. Elution fractions were collected and analysed using SDS-PAGE.

**Supplementary methods 2 - Static Light Scattering**

To investigate whether Tpp49Aa1 was present as a monomer in solution we used static light scattering (RALS) and refractive index (RI) measurements. The Zetasizer MicroV system (Malvern Instruments Ltd., Malvern, UK) was used to quantify the size distribution of Tpp49Aa1 particles based upon time-dependent fluctuations in the scattered light intensity (at 90° scattering angle), due to the Brownian motion of the protein in solution. Concentration was determined by a refractive index (RI) detector (VE 3580, Viscotek Corp). Protein samples (100 µL at 1 mg/ mL) were prepared and solubilised as above, with protein-containing fractions pooled and concentrated from the first round of SEC (supplementary methods 1). The sample was again separated via SEC using a Superdex 75 Increase 10/300 GL with a flow rate of 0.8 mL/min, coupled to the Zetasizer and the refractive index (RI) detector. Eluted samples were measured every ~3 s at 30°C. Data were collected and analysed in OmniSEC software (Ver 5.12) and calibrated to BSA (1 mg/ mL).

**Supplementary methods 3 – Molecular Dynamics Simulations**

To investigate the structural stability of Cry48Aa1-Tpp49Aa1 models generated, 100 ns molecular dynamics (MD) simulations were performed using GROMACS (v.2020.1) (1). Models were solvated using a 3-point model and neutralised within a cubic box. The AMBER99SB force field was applied (2). For energy minimisation, the steepest descent algorithm (step sizes = 0.01, maximum number of steps = 50,000) was used. Equilibration steps were performed for 100 ps and involved using an isothermal-isochoric ensemble and an isothermal-isobaric ensemble to stabilize the temperature and pressure of the system respectively. Pressure coupling was achieved using a Parrinello-Rahman barostat.

Simulations of 100 ns were performed at a constant temperature of 300 K using a velocity rescaling thermostat. Separate couplings were applied for proteins and non-proteins. A constant pressure of 1.0 bar was achieved using a Parrinello-Rahman barostat with an isothermal compressibility of 4.5 x 10^-5^ bar^-1^. Pressure was coupled isotropically using a coupling constant of 2.0 ps. A timestep of 2.0 fs was used to integrate Newton’s equations of motion. Non-bonded long-range electrostatic interactions were calculated using the Particle-Mesh Ewald method and a 10 Å cut-off. A 10 Å cut-off was also applied for van der Waals interactions. The LINCS constraint algorithm was used to constrain bonded hydrogen atoms.
